# Sex differences in hippocampal cytokine networks after systemic immune challenge

**DOI:** 10.1101/378257

**Authors:** Julie E. Finnell, Ian C. Speirs, Natalie C. Tronson

## Abstract

Increased production of cytokines in the in the brain during illness or injury modulates physiological processes, behavior, and cognitive function. It is likely that the pattern of cytokines, rather than the activation of any individual cytokine, determines the functional outcome of neuroimmune signaling. Cytokine networks may thus be particularly useful for understanding sex differences in immune and neuroimmune activation and outcomes. In this project, we aimed to determine the activation and resolution of hippocampal cytokine networks in both male and female mice. We measured 32 cytokines in the hippocampus and periphery of male and female mice at rest, 2, 6, 24, 48, and 168 hours after an acute systemic injection of lipopolysaccharide (LPS; 250μg/kg). We hypothesized that males and females would exhibit both differences in individual cytokine levels and differences in network dynamics of hippocampal cytokines. Cytokines with sex-specific activation by LPS included male-specific elevations of IFNɣ, CSF1, CSF2, and IL-10; and female-specific activation of the IL-2 family and IL-4. We also observed differences in time course, where females showed more rapid elevations, and faster resolution of cytokine activity compared with males. Network analysis using ARACNE and Cytoscape demonstrated markedly different hippocampal cytokine networks across sex even at baseline, and sex differences in cytokine network activation states in response to LPS. Analysis of global shifts in cytokine concentrations further identified a period of cytokine and chemokine downregulation at 48 hours that was more pronounced in females compared with males. Together, these findings demonstrate that sex differences in neuroimmune responses include both differences in intensity of the cytokine response, and importantly differences in cytokine networks activated. Such sex differences in cytokine networks in the brain are likely critical for short and long-term functional outcomes associated with neuroimmune activation.

## 1. INTRODUCTION

There are clear sex differences in the peripheral immune response ^1–6^ and in the physiological and behavioral responses to an immune challenge ^7–11^. Sex differences in immune responses are both quantitative—women and female rodents show stronger febrile responses, and greater regulation of mood ^9,12–14^; and qualitative— men and women showing different kinds of responses, including male-specific “sighing” ^15^ and female-specific changes in pain sensitivity ^8^. These sex differences in behavioral and physiological responses to peripheral immune challenge are likely driven by differences in the cytokines activated following an immune activation. For example, Tonelli and colleagues ^12^ observed that female rats showed greater Interleukin (IL)-6 and Tumor Necrosis Factor (TNF) α gene expression in the hippocampus 24 hours after lipopolysaccharides (LPS) injection. In contrast, males show more IL-1β after LPS ^14^ and a stronger inflammatory response in the hippocampus after stress ^5,16,17^. Cytokines also play different roles in the brain of males and females. Hippocampal IL-2 impairs neurogenesis only in males ^18^, whereas IL-13 mediates symptoms in models of multiple sclerosis only in females ^19^. Males and females therefore show different functional correlates of neuroimmune signaling, perhaps determined by sex-specific networks of cytokine activation in the brain.

As in the periphery, IL-1β, IL-6, and TNFα have been particularly well studied for their functional correlates in the brain after many different inflammatory challenges ^20–29^. Despite the consistent activation of these cytokines, different immune challenges lead to unique functional correlates ^30,31^. As such, the precise role of individual cytokines likely depends on the exact pattern of immune cells and cytokines activated concurrently ^31–33^. Immune signaling exhibits complex and tightly regulated expansion and resolution of cytokines, a process that is well described in the periphery ^33–36^. Here, after the initial challenge, acute phase cytokines TNF and IL-1 rapidly accumulate and stimulate the production of other cytokines such as IL-6 and interferon ɣ (IFNɣ). These, in turn, recruit specific immune cells, and release additional cytokines and chemokines, including the regulatory cytokines IL-10, IL-4, and IL-13 ^34^. The precise pattern of cytokine activation and resolution is dependent on several factors: the stimulant, which acute phase cytokines are induced, which immune cells are recruited, what tissue the response is in, and the concurrent endocrine and inflammatory environment ^30,33,37^. Such network-dependent patterns of immune signaling means that in the periphery ^33,37–39^— and in the brain ^31,40–42^— functional outcomes are determined by both the magnitude of activation and the specific pattern of cytokines induced. Therefore, differences in the cytokine networks activated in the brain in females compared with males are likely critical determinants of sex-specific outcomes after an immune challenge.

In this project, we identified sex-specific patterns of cytokine activation in the hippocampus after a systemic LPS challenge, despite broad similarities in peripheral cytokine patterns. We examined 32 cytokines in the hippocampus and serum of males and females at rest and from 2 hours to 7 days after an acute, systemic LPS injection. These cytokine and chemokine data were subsequently used in conjunction with ARACNE mutual information analysis and Cytoscape to statistically model cytokine networks at each time point. Cytokine networks are of particular importance as these statistical maps provide key information about the cell types involved, and point to the general function of cytokines at any given point in time. We anticipated that both males and females would show increased cytokine signaling, but that patterns of cytokine activation, kinetics, magnitude of changes, and functional cytokine networks would differ between the sexes. Delineating the patterns of cytokine activation in the brain after a systemic immune challenge in males and females is essential for understanding how cytokine-dependent signaling may modulate behavior and vulnerability in a sex-specific manner.

## 2. METHODS

### 2.1 Animals

38 male and 40 female C57Bl6 mice (Harlan Laboratories, Indianapolis, IN), 9-11 weeks old were allowed to habituate to the colony room for at least 5 days prior to injections. Animals were housed individually in standard mouse caging, with *ad lib* access to food and water, a 12:12h light:dark cycle (lights on 7am-7pm) and temperature maintained at 70°F. Individual housing of adult males is required to reduce fighting-induced stress, and is consistent with AAALAC guidelines on management of fighting in mice ^43^. To keep social influences constant across animals of both sexes, all animals, regardless of sex, were maintained in individual housing. Due to independent social structures of both male and female mice ^44,45^, individual housing is ecologically appropriate for both sexes and does not increase variance in either sex ^46^. All procedures were approved by University of Michigan Committee for the Care and Use of Animals.

### 2.2 Lipopolysaccharide Injections

Mice were given single intraperitoneal (i.p.) injections of lipopolysaccharide (LPS; E coli, O111:B4; Sigma, St Louis) (250μg/kg in 2mL/kg) or sterile phosphate buffered saline (2mL/kg; PBS). This dose of LPS has previously been demonstrated to result in a robust cytokine response in the hippocampus ^47^ and memory impairments ^48,49^ in adult male mice. All animals were monitored after injection for recovery from LPS induced illness.

### 2.3 Tissue Dissections

0 (males n=13; females n=13), 2 (males n=5; females n=5), 6 (males n=5; females n=5), 24 (males n=5; females n=5), 48 (males n=5; females n=5), or 168 (males n=5; females n=5) hours after LPS or PBS (males n=13; females n=13) injection, mice were rapidly decapitated, hippocampi immediately dissected and frozen in liquid nitrogen. Trunk blood was collected and allowed to clot at room temperature for approximately 30 minutes, then centrifuged at 1000 rpm for 20 minutes and serum without red blood cells was collected.

### 2.4 Tissue Preparation

Hippocampal tissue was sonicated in low-detergent RIPA buffer (0.01% Triton-X in 1X PBS, with NaVO4, NaF, EDTA, and protease and phosphatase inhibitor (HALT)), centrifuged at 11500rpm, and supernatant collected and stored at -20°C. Bradford assays (Bio-rad Laboratories, Hercules, CA) were used to determine protein concentration of each sample.

### 2.5 Cytokine Assays

32-plex Magplex cytokine assays (EMD Millipore Darmstadt, Germany) ^50^ were used according to the protocols provided, with minor adaptations for brain tissue. Briefly, 25μL of diluted serum were loaded per well. Hippocampal tissue samples were filtered and diluted to concentrations of 1μg/μL prior to loading 25μL of per well. All samples were run in duplicate, and all plates were read on a MagPix (Luminex Corp, Austin TX) machine.

The following cytokines were assayed: CSF1, CSF2, CSF3, IL-1α, IL-1β, IL-2, IL-3, IL-4, IL-5, IL-6, IL-9, IL-10, IL-12p40, IL-12p70, IL-13, IL-15, IL-17, IFNɣ, CXCL1, CXCL2, CXCL5, CXCL9, CXCL10, CCL2, CCL3, CCL4, CCL5, CCL11, TNFα, VEGF, LIF, and LIX. Wash was run between each well to prevent build-up of beads and tissue on the plate. Cytokine concentrations were calculated as pg/mg of hippocampal tissue and pg/mL of serum via the native Luminex software.

### 2.6 Cytokine network analysis

Cytoscape and the ARACNE algorithm were used to generate cytokine networks from collected Magplex data. Unlike Bayesian networks that rely upon Pearson’s correlational analysis^51^, ARACNE relies upon mutual information (MI) analysis to determine the relatedness of two or more biological factors (cytokines, genes, etc.). This is a major advantage as ARACNE does not make assumptions about the nature of the relatedness of two factors and is capable of detecting a broader range of interactions that do not strictly adhere to a 1:1 linear relationship. Originally developed by Margolin et al. ^52,53^, ARACNE determines MI between two factors (X and Y) by calculating the entropy (H) of each factor and the factors in combination using the following mathematical expression:

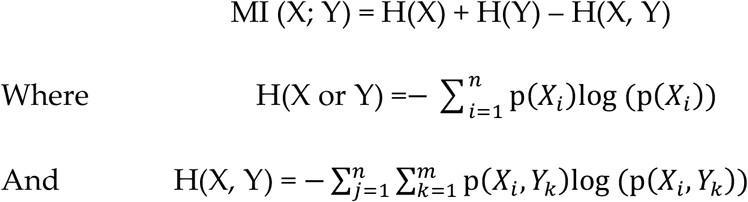

The ARACNE algorithm calculates MI scores for each cytokine and chemokine combination in the supplied data set and then statistically compares MI values to “0”, which represents true mathematical independence. Resulting p-values are then used to generate cytokine networks in Cytoscape. Importantly, ARACNE and Cytoscape use p-value thresholds in the generation of their networks. As such, all connections shown for a given network fall within the user defined p-value threshold. For the current study, network stability was tested between p = 0.0001 – 0.07. Final significance thresholds for each network are reported in the results below.

The strength of individual connections (network edges) between cytokines and chemokines (network nodes) were visualized using continuous mapping of MI scores ranging from 0.5 pt to 3 pt thickness: Hippocampus minimum MI = 0.2, maximum MI = 0.8; Serum – minimum MI = 0.2, maximum MI = 1. A summary table of MI scores are shown in Supplemental Tables 1-6. Cytokine concentrations at 2, 6, 24, 48, and 168 hours post LPS injection were transformed to log_2_(fold) from saline controls which were used for continuous mapping of relative shifts in cytokine concentration. Additionally, average log_2_(fold) was calculated and compared across time points in males and females to determine whether sex contributed to global differences in cytokine concentrations.

### 2.7 Statistical Analysis

Statistical analyses were conducted on protein concentration data. PBS treated animals were collapsed into one group for each sex. Two-way multivariate ANOVA was used, with Sex and Time post injection as factors. Effect size was estimated using partial Eta-squared (η^2^p). Post-hoc tests were used to further examine significant effects, with Bonferroni corrections for multiple comparisons. To determine whether males and females differed at baseline, we compared baseline data from both sexes run on the same multiplex plate. As no differences were found, we directly compared changes in individual cytokines between males and females, without the small variations in baseline, we normalized all values to the 0 timepoint (PBS) controls for each sex and compared changes from baseline using these values. Additionally, Wilcoxon signed rank tests were applied to determine whether global shifts in cytokine concentrations, as determined by increases or decreases in log_2_(fold), were significantly different from resting saline controls (log_2_(fold)=0). Analyses were carried out using SPSS and GraphPad Prism, and all data is expressed as mean ± SEM.

## 3. RESULTS

### 3.1 Cytokines in the hippocampus

Of 32 cytokines tested, 27 exhibited a significant elevation in the hippocampus after systemic LPS injection (Main effect of TIME, Table 1), four (LIF, IL-3, IL-5, and IL-15) were not significantly elevated when collapsed across males and females, and CXCL5 (LIX) was excluded as it never reached minimum detectable concentration as defined by the Magplex kit. Post-hoc tests demonstrated rapid activation of IL-1α (*p* < 0.01), IL-2 (*p* < 0.001), IL-4 (*p* < 0.01), IL-6 (*p* < 0.001), IL-9 (*p* < 0.001), IL-12p40 (*p* < 0.01), IL-13 (*p* < 0.05), IL-17 (*p* < 0.001), CSF3 (*p* < 0.001), CXCL1 (*p* < 0.001), CXCL2 (*p* < 0.001), CCL11 (*p* < 0.001) and VEGF (*p* < 0.001) 2 hours after systemic LPS injection. Of these, only IL-4 and CCL11 showed fast resolution with no ongoing increase at the 6-hour timepoint. Activation of CSF1 (*p* < 0.001), CSF2 (*p* < 0.001), TNFα (*p* < 0.001), IFNγ (*p* < 0.001), IL-1β (*p* < 0.001), IL-7 (*p* < 0.05), IL-10 (*p* < 0.001), IL-12p70 (*p* < 0.01), CXCL9 (*p* < 0.001), CXCL10 (*p* < 0.001), CCL2 (*p* < 0.001), and CCL5(*p* < 0.001) was first observed 6 hours after LPS. Elevated levels of several cytokines, including CSF2 (*p* < 0.01), TNFα (*p* < 0.05), IFNγ (*p* < 0.001), IL-6 (*p* < 0.001), IL-9 (*p* < 0.01), IL-10 (*p* < 0.01), IL-12p40 (*p* < 0.05), CXCL1 (*p* < 0.05), CXCL9 (*p* < 0.001), CXCL10 (*p* < 0.001), and CCL2 (*p* < 0.001) persisted at least 24 hours after LPS injection. There were no ongoing elevations in cytokine levels 48 hours after LPS.

**Table 1.** Hippocampal cytokines: Statistical analysis summary. Overall multivariate ANOVA results for the Main effect of TIME and the interaction effect (SEX x TIME). \**p* < 0.05, \*\**p* < 0.01, \*\*\**p* < 0.001.

| CYTOKINE | MAIN EFFECT: TIME | INTERACTION: SEX x TIME |
| --- | --- | --- |
| **Hippocampal cytokines differentially activated in males and females** |  |  |
| IL-1α | *** $F_{(5,66)} = 13.09$ $\eta^2_p = 0.50$ | *** $F_{(5,66)} = 9.11$ $\eta^2_p = 0.41$ |
| IL-1β | *** $F_{(5,66)} = 6.93$ $\eta^2_p = 0.34$ | *** $F_{(5,66)} = 8.31$ $\eta^2_p = 0.39$ |
| TNFα | *** $F_{(5,66)} = 12.46$ $\eta^2_p = 0.486$ | *** $F_{(5,66)} = 6.19$ $\eta^2_p = 0.32$ |
| IL-6 | *** $F_{(5,66)} = 30.19$ $\eta^2_p = 0.70$ | ** $F_{(5,66)} = 3.62$ $\eta^2_p = 0.22$ |
| IL-10 | *** $F_{(5,66)} = 20.21$ $\eta^2_p = 0.60$ | *** $F_{(5,66)} = 5.48$ $\eta^2_p = 0.29$ |
| IL-4 | *** $F_{(5,66)} = 13.83$ $\eta^2_p = 0.51$ | *** $F_{(5,66)} = 5.33$ $\eta^2_p = 0.29$ |
| IL-13 | *** $F_{(5,66)} = 23.28$ $\eta^2_p = 0.64$ | *** $F_{(5,66)} = 2.12$ $\eta^2_p = 0.42$ |
| IL-2 | *** $F_{(5,66)} = 13.88$ $\eta^2_p = 0.51$ | ** $F_{(5,66)} = 4.15$ $\eta^2_p = 0.24$ |
| IL-15 | n.s. | *** $F_{(5,66)} = 6.23$ $\eta^2_p = 0.32$ |
| IFNγ | *** $F_{(5,66)} = 33.36$ $\eta^2_p = 0.72$ | *** $F_{(5,66)} = 27.37$ $\eta^2_p = 0.68$ |
| CXCL9 (MIG) | *** $F_{(5,66)} = 40.47$ $\eta^2_p = 0.75$ | *** $F_{(5,66)} = 7.50$ $\eta^2_p = 0.36$ |
| CXCL10 (IP-10) | *** $F_{(5,66)} = 115.95$ $\eta^2_p = 0.90$ | *** $F_{(5,66)} = 5.62$ $\eta^2_p = 0.30$ |
| CSF1 (M-CSF) | *** $F_{(5,66)} = 17.49$ $\eta^2_p = 0.57$ | *** $F_{(5,66)} = 9.40$ $\eta^2_p = 0.42$ |
| CSF2 (GM-CSF) | *** $F_{(5,66)} = 16.41$ $\eta^2_p = 0.55$ | ** $F_{(5,66)} = 4.78$ $\eta^2_p = 0.27$ |
| CSF3 (G-CSF) | *** $F_{(5,66)} = 81.25$ $\eta^2_p = 0.86$ | *** $F_{(5,66)} = 27.57$ $\eta^2_p = 0.68$ |
| CXCL1 (KC) | *** $F_{(5,66)} = 103.11$ $\eta^2_p = .89$ | *** $F_{(5,66)} = 25.50$ $\eta^2_p = 0.66$ |
| CCL3 (MIP1α) | * $F_{(5,66)} = 2.62$ $\eta^2_p = 0.17$ | *** $F_{(5,66)} = 19.01$ $\eta^2_p = 0.59$ |
| CCL5 (RANTES) | *** $F_{(5,66)} = 8.97$ $\eta^2_p = 0.41$ | *** $F_{(5,66)} = 13.09$ $\eta^2_p = 0.50$ |
| IL-12p40 | *** $F_{(5,66)} = 16.12$ $\eta^2_p = 0.55$ | ** $F_{(5,66)} = 3.94$ $\eta^2_p = 0.23$ |
| VEGF | *** $F_{(5,66)} = 15.22$ $\eta^2_p = 0.54$ | * $F_{(5,66)} = 2.34$ $\eta^2_p = 0.15$ |
| IL-3 | n.s. | * $F_{(5,66)} = 2.81$ $\eta^2_p = 0.18$ |
| **Hippocampal cytokines activated in both sexes** |  |  |
| IL-9 | *** $F_{(5,66)} = 16.52$ $\eta^2_p = 0.56$ | n.s. |
| CCL2 (MCP1) | *** $F_{(5,66)} = 31.92$ $\eta^2_p = 0.71$ | n.s. |
| CCL11 (eotaxin) | *** $F_{(5,66)} = 17.06$ $\eta^2_p = 0.56$ | n.s. |
| CCL4 (MIP1β) | *** $F_{(5,66)} = 6.12$ $\eta^2_p = 0.32$ | n.s. |
| CXCL2 (MIP2α) | *** $F_{(5,66)} = 17.93$ $\eta^2_p = 0.58$ | n.s. |
| IL-12p70 | *** $F_{(5,66)} = 8.24$ $\eta^2_p = 0.38$ | n.s. |
| IL-17 | *** $F_{(5,66)} = 17.12$ $\eta^2_p = 0.57$ | n.s. |
| IL-7 | ** $F_{(5,66)} = 4.15$ $\eta^2_p = 0.24$ | n.s. |

#### 3.1.2 Sex differences in activation

Both males and females exhibited a robust neuroimmune response in the hippocampus, and sex differences were observed in specific cytokines activated, the time course of cytokine response, and the magnitude of activation. In the hippocampus, 20 of 31 cytokines showed differential activation in males and females, via a significant SEX x TIME interaction (see Table 1). IL-7, IL-9, IL-12p70, IL-17, CCL2, CCL4, CCL11, CXCL2 and CXCL5 showed no significant SEX x TIME interaction, but were elevated in the hippocampus in both males and females, whereas IL-5, and LIF were not elevated in either sex. IL-3 and VEGF showed very small effect size for SEX x TIME interaction (η^2^p = 0.18; η^2^p = 0.15, respectively). Post-hoc tests were used to determine significant elevations at each timepoint. No significant elevations in males or females were observed at 168 hours. Finally, to determine whether elevations in cytokine level differed between sex at each time point for cytokines that showed significant Sex x Time effects, we used post-hoc tests for data normalized to baseline controls.

##### 3.1.2.1 IL-1, TNFα, and IL-6

IL-1α and IL-1β were elevated in the hippocampus after systemic LPS. IL-1β was significantly increased in both males and females, with different time courses [SEX x TIME: *F*(5,66) = 8.31, *p* < 0.001]. Females showed rapid activation 2 hours after LPS injection (*p* < 0.05; m *vs* f: *p* < 0.05), which was resolved by 6 hours (*p* = 0.20). In contrast males showed a later increase, peaking at 6 hours after injection (*p* < 0.01; 24h: *p* = 0.39) (Fig. 1a). IL-1α showed a female-specific increase in the hippocampus 6 and 24 hours after LPS, and resolved by 48 hours [SEX x TIME: *F*(5,66) = 9.11; female 6h: *p* < 0.001; 24h: *p* < 0.01; male 6h: *p* = 0.67 24h: *p* = 0.65; m *vs* f 6h: *p* < 0.001; 24h: *p* < 0.01].

**Figure 1.**
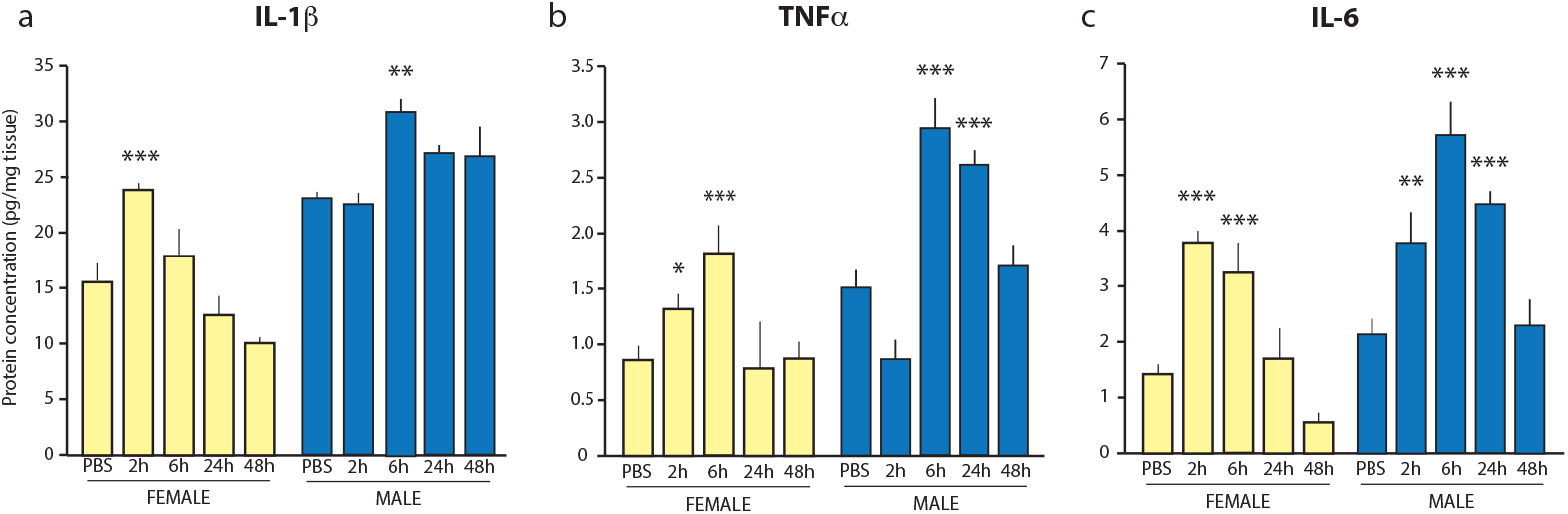
IL-1β, TNFα, and IL-6 are increased in the hippocampus of males and females after systemic LPS injection. (a) IL-1β is increased 6 hours after LPS injection in males and 2 hours after LPS in females. (b) TNFα is elevated at 6 hours after LPS in both male and female hippocampus. In males, this activation persists for at least 24 hours. (c) IL-6 is increased 6-24 hours after LPS in males, and earlier in females (2-6 hours). \**p* < 0.05, \*\**p* < 0.01, *** *p* < 0.001 all *cf* PBS.

TNFα showed significant differences in time course of activation in males compared with females [SEX x TIME: *F*(5,66) = 6.19, *p* < 0.001;m *vs* f: *p* < 0.01]. Both sexes showed peak levels at 6 hours (Female: *p* < 0.01; Male: *p* < 0.001; m *vs* f: *p* =0.18), however this response persisted at least 24 hours only in male mice (Male: *p* < 0.01; Female: *p* = 1.00; m *vs* f: *p* < 0.05) (Fig. 1b).

IL-6, like IL-1β, exhibited significant increases in the hippocampi of both male and female mice, with sex differences in time course [SEX x TIME: *F*(5,66) = 3.62,*p* < 0.01]. Again, females mounted earlier responses, with increased IL-6 observed at 2-6 hours post injection, that had returned to baseline by 24 hours (2h: *p* < 0.001; 6h: *p* < 0.01; 24h: *p* = 0.853; 2h m *vs* f: *p* < 0.001). Males showed elevations at 6 hours that persisted for at least 24 hours (2h: *p* = 0.100; 6h *p* < 0.001; 24h: *p* < 0.01) (Fig. 1c).

##### 3.1.2.2 IL-10, IL-4, IL-13

We also observed striking sex differences in IL-10, IL-4, and IL-13 (Fig. 2). Males showed significant activation of IL-10 6 and 24 hours after LPS injection [*F*(5,66) = 5.48, *p* < 0.001; Males 6h: *p* < 0.001, 24h: *p* < 0.01], whereas females showed only a smaller elevation at 6h (Females 6h: *p* < 0.01; 24h *p* = 0.36; m *vs* f 6h: *p* < 0.01; 24h: *p* < 0.01) (Fig. 2a). In contrast, IL-4 was increased only in females [SEX x TIME: *F*(5,66) = 5.33, *p* < 0.001; Females 2h: *p* < 0.001; Males: *p* = 0.21; m *vs* f: *p* < 0.001] (Fig. 2b). Finally IL-13, an “IL-4-like cytokine” was differentially activated in females and males over time [TIME: *F*(5,66) = 16.52, *p* < 0.001; SEX x Time *F*(5,66) = 2.12, *p* < 0.001]. IL-13, like IL-6 and IL-1β, was induced earlier in females, peaked at 2 hours and returned to baseline by 6 hours post LPS (2h: *p* < 0.01; 6h: *p* = 0.88; m *vs* f 2h: *p* < 0.01), whereas in males, IL-13 levels increased later, and stayed elevated for longer (6h: *p* < 0.00; 24h *p* = 0.018; m *vs* f 6h: *p* < 0.01; 24h *p* < 0.001) (Fig. 2c).

**Figure 2.**
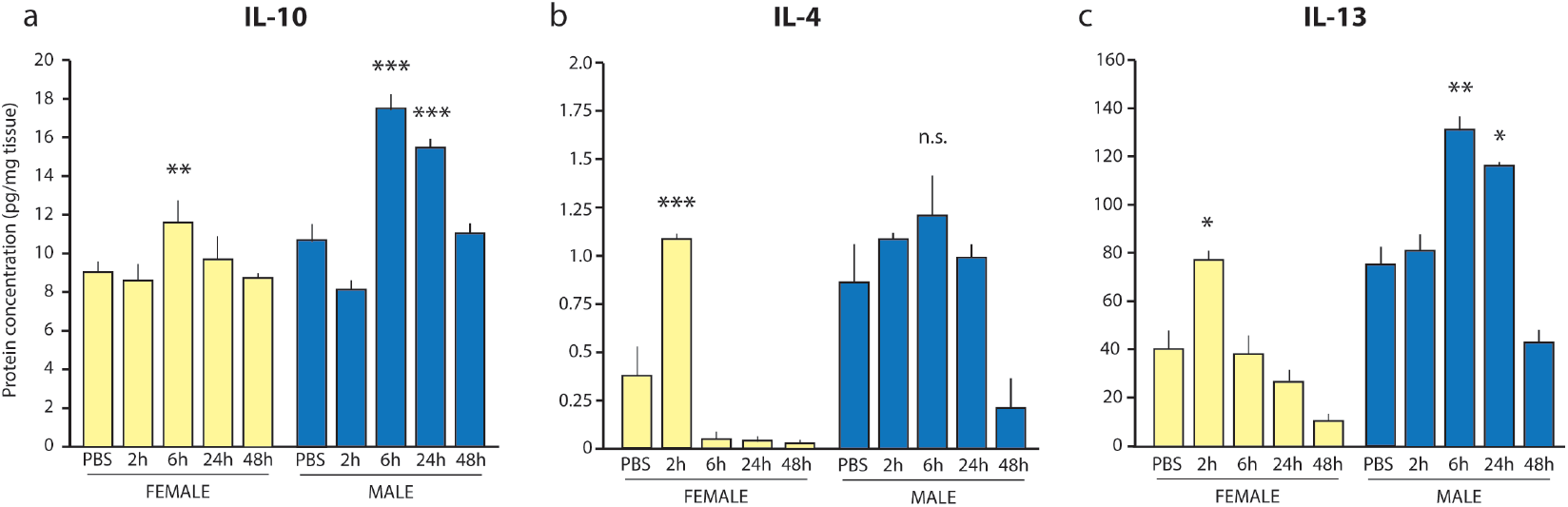
Regulatory cytokines IL-10, IL-4, and IL-13 are differentially activated in males and females. (a) IL-10 is elevated only in males 6-24 hours after LPS injection; (b) IL-4 is elevated only in females 2 hours after LPS injection; (c) IL-13 is activated in females at 2 hours, and in males 6-24 hours after LPS. \**p* < 0.05, \*\**p* < 0.01, \*\*\**p* < 0.001 all cf PBS.

##### 3.1.2.3 IL-2 family cytokines (IL-2, IL-15, IL-9)

The IL-2 family of cytokines was also activated in both sexes, with a bias towards activation of IL-2 in male animals and IL-15 in females (Fig. 3). In males and females, IL-2 was elevated 6 hours after LPS injection, and but this persisted only in males [SEX x TIME: *F*(5,66) = 4.15, *p* <0.01; Females 6h: *p* < 0.05; 24h: *p* < 0.383; Males 6h: p < 0.01; 24h p < 0.05; 48h *p* < 0.01; m *vs* f 6h: *p* = 0.14; 48h: *p* < 0.05] (Fig. 3a). In contrast, IL-15 was elevated at 2 hours [SEX x TIME: *F*(5,66) = 6.23, *p* < 0.001; Females 2h: *p* < 0.01; Males 2h: *p* = 0.24; m *vs* f 2h: *p* <0.001] (Fig. 3b). IL-9 was significantly increased in both males and females at 6, 24, and 48 hours after LPS [TIME: *F*(5,66) = 16.52, *p* < 0.001; SEX x Time *F*(5,66) = 2.12, *p* > 0.05].

**Figure 3.**
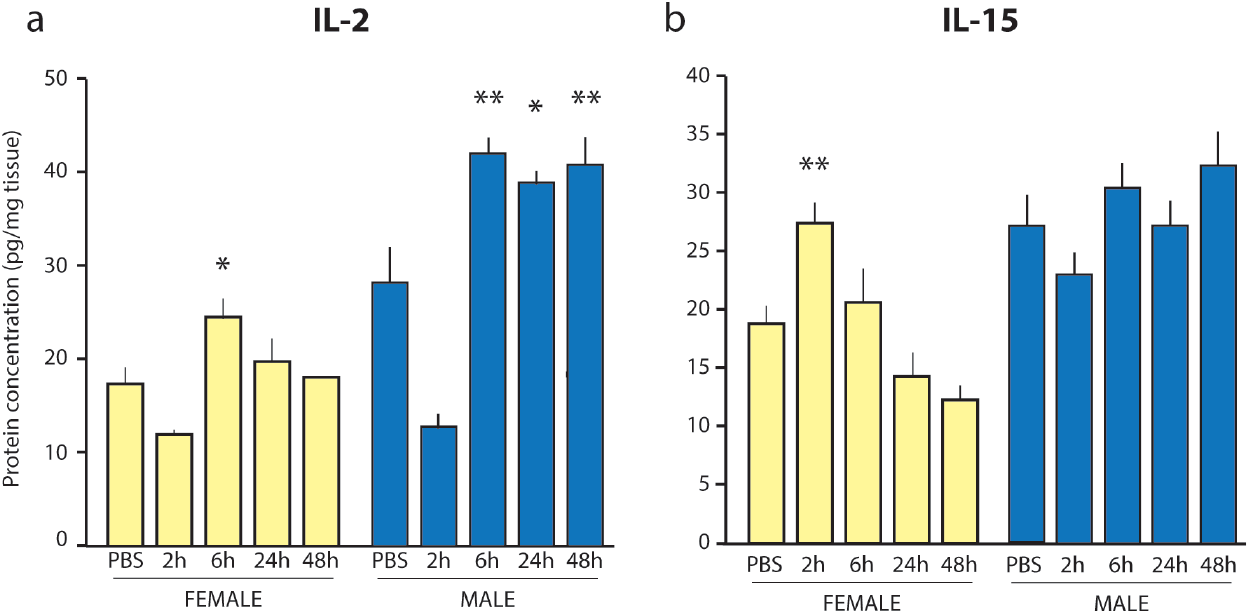
IL-2 and IL-15 are increased only in females. Both (a) IL-2 is elevated at 6 hours and (b) IL-15 is elevated 2 hours after LPS in females, but not males. \**p* < 0.05, \*\**p* < 0.01 all cf PBS.

##### 3.1.2.4 IFNɣ, CXCL9, and CXCL10

IFNɣ showed male-specific activation in the hippocampus [SEX x TIME *F*(5,66) = 27.37, *p* < 0.001]. Males, but not females, showed elevations at 6 hours and 24 hours (Males 6h: *p* < 0.001; 24h *p* < 0.001) after LPS injection (Fig. 4a).

**Figure 4.**
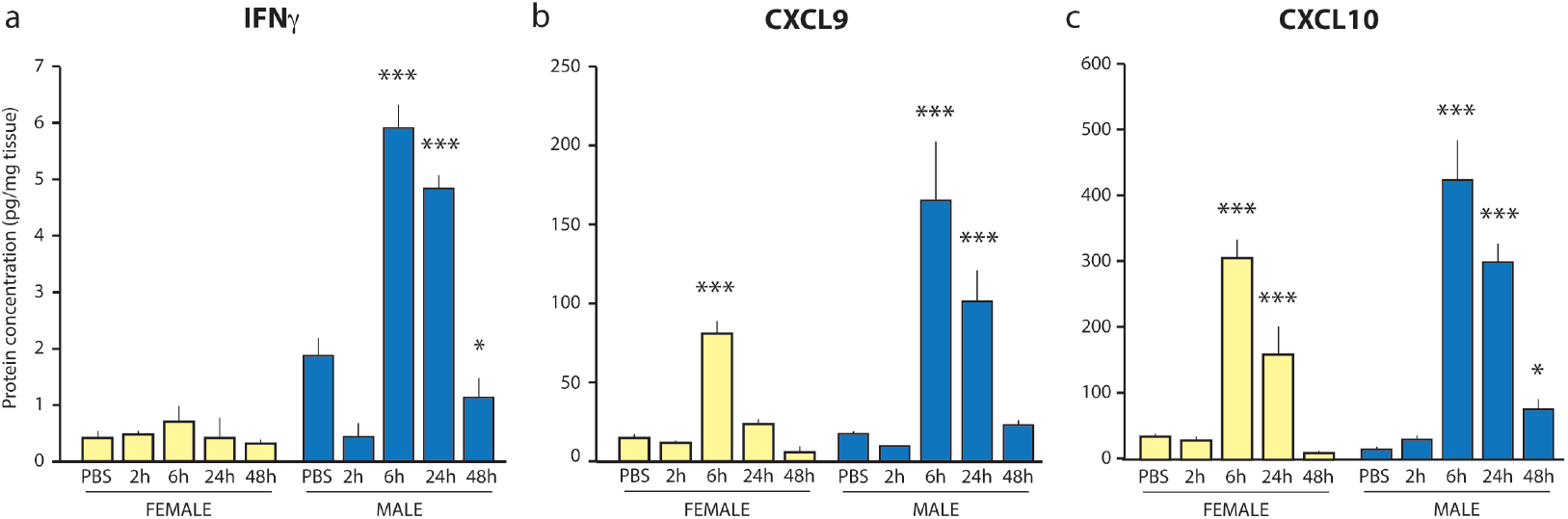
IFNγ, CXCL9, and CXCL10. (a) IFNγ is elevated only in males 6-24 hours after systemic LPS injection. (b) CXCL9 is elevated in both males and females 6 hours after LPS, and persists for at least 24 hours only in males. (c) CXCL10 is increased in both males and females 6-24 hours after LPS. \**p* < 0.05, \*\**p* < 0.01, \*\*\**p* < 0.001 all *cf* PBS.

The CXCR3 ligands CXCL9 (MIG) and CXCL10 (IFN gamma induced protein 10; IP-10) were increased in both males and females [CXCL9 TIME: *F*(5,66) = 40.47, *p* < 0.001;CXCL10 TIME: *F*(5,66) = 115.95, *p* < 0.001]. Both CXCL9 and 10 showed sex differences in activation with higher elevations in males compared to females [SEX x TIME: *F(*5,66) = 7.50, *p* < 0.001; CXCL10 SEX x TIME: *F*(5,66) = 5.62, *p* < 0.001]. CXCL9 exhibited pronounced differences in time course of activation between males and females, with females showing activation of CXCL9 only at 6 hours (6h: *p* < 0.001; 24h: *p* = 0.33), whereas CXCL9 persisted for at least 24 hours in males (6h: *p* < 0.001; 24h: *p* < 0.01; 48h: *p* = 0.50) (Fig. 4b). CXCL10 was elevated in both sexes at 6h and 24h (all *p* < 0.001) and persisted in males at 48h (Males: *p* < 0.05; Females *p* = 0.91), nevertheless, CXCL10 showed greater elevations than females at all time points (m *vs* f 6h: *p* < 0.001; 24h: *p* < 0.001; 48h: *p* < 0.05) (Fig. 4c).

##### 3.1.2.5 Colony Stimulation Factor (CSF1, CSF2, CSF3)

All three CSF family members show significant elevation after LPS and differential activation in males and females [CSF1 TIME: *F*(5,66) = 17.59, *p* < 0.001; SEX x TIME: *F*(5,66) = 9.40, *p* < 0.001; CSF2 TIME: *F*(5,66) = 16.41, *p* < 0.001; SEX x TIME: *F*(5,66) = 4.78, *p* < 0.001; CSF3 TIME: *F*(5,66) = 81.25; *p* < 0.001] (Fig. 5).

**Figure 5.**
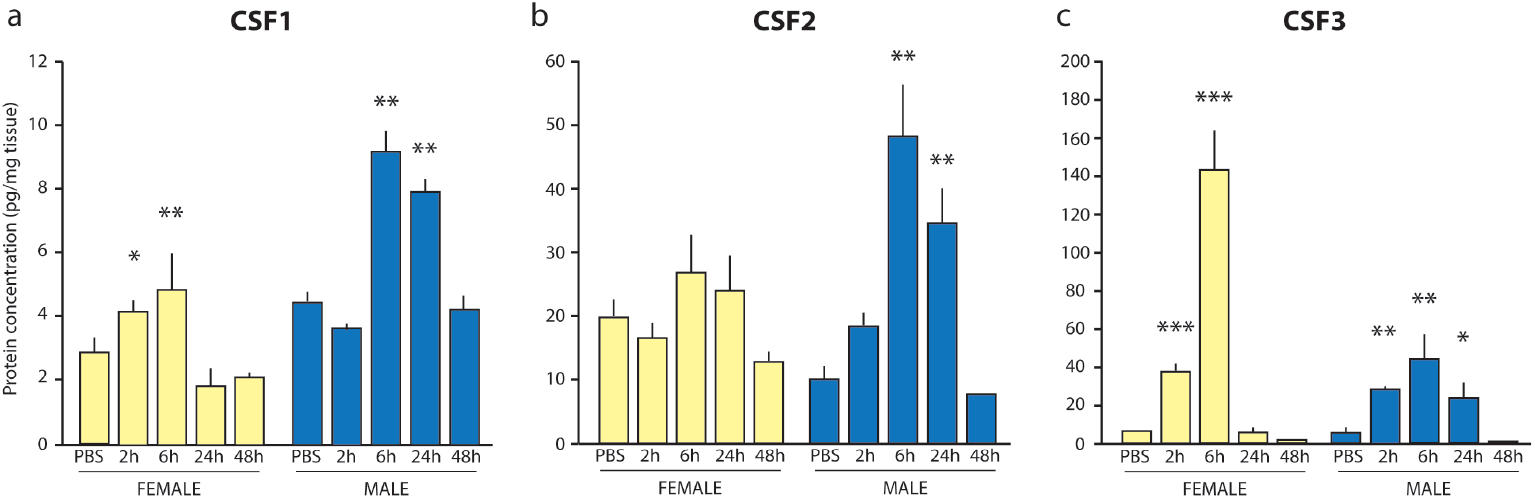
Colony Stimulating Factor Family. (a) CSF1 and (b) CSF2 are elevated in males but not females, 6-24 hours after LPS injection. (c) CSF3 is significantly elevated in females 2-6 hours after LPS, and to a lesser degree in males at the 6 hour timepoint. \**p* < 0.05, \*\**p* < 0.01, \*\*\**p* < 0.001 all *cf* PBS.

CSF1 and CSF2 showed male-specific activation, and were significantly elevated 6 and 24 hours after LPS injection [CSF1 2h: *p* = 0.972, 6h: *p* < 0.001, 24h: *p* < 0.001; CSF2 2h: *p* = 0.78, 6h: *p* < 0.001; 24h: *p* < 0.005], whereas females showed no change in either CSF1 or CSF2 at any time point (all *p* > 0.1) (Fig. 5a,b). In contrast, females showed elevations in CSF3 at both 2 and 6 hours after LPS injection [2h: *p* = 0.004; 6h; *p* < 0.001; 24h: *p* = 0.994], and males show an increase in CSF3 only at 6h post LPS [2h: *p* = 0.127; 6h; *p* < 0.001; 24h: *p* = 0.223], although to a lesser degree than in females (m vs f: *p* < 0.001) (Fig. 5c).

##### 3.1.2.6 CXCR2 Ligands (CXCL1 and CXCL2, Gro family)

CXCL1 and CXCL2 were strongly activated in the hippocampus in both males and females after LPS injection (Fig. 6). CXCL1 showed significant sex differences in activation [SEX x TIME *F*(5,66) = 25.50, *p* < 0.001]. Both males and females showed significant elevations at 2 hours [all *p* < 0.001] and 6 hours (all *p* < 0.001) but only males showed persisted elevations 24 hours after LPS [Males: *p* < 0.01; Females: *p* = 0.11]; nevertheless, females showed significantly greater activation compared with males at 2 and 6 hours after LPS (m *vs* f 2h: *p* < 0.001; 6h: *p* < 0.001) (Fig. 6a). CXCL2 was significantly elevated 2 and 6 hours after LPS with no difference between sexes [TIME: *F*(5,66) = 17.93, *p* < 0.001; SEX x TIME: *F*(5,66) = 1.489, *p* = 0.205; Males 2h: *p* < 0.01; 6h: *p* < 0.01; Females 2h: *p* < 0.001; 6h: *p* < 0.001] (Fig. 6b).

**Figure 6.**
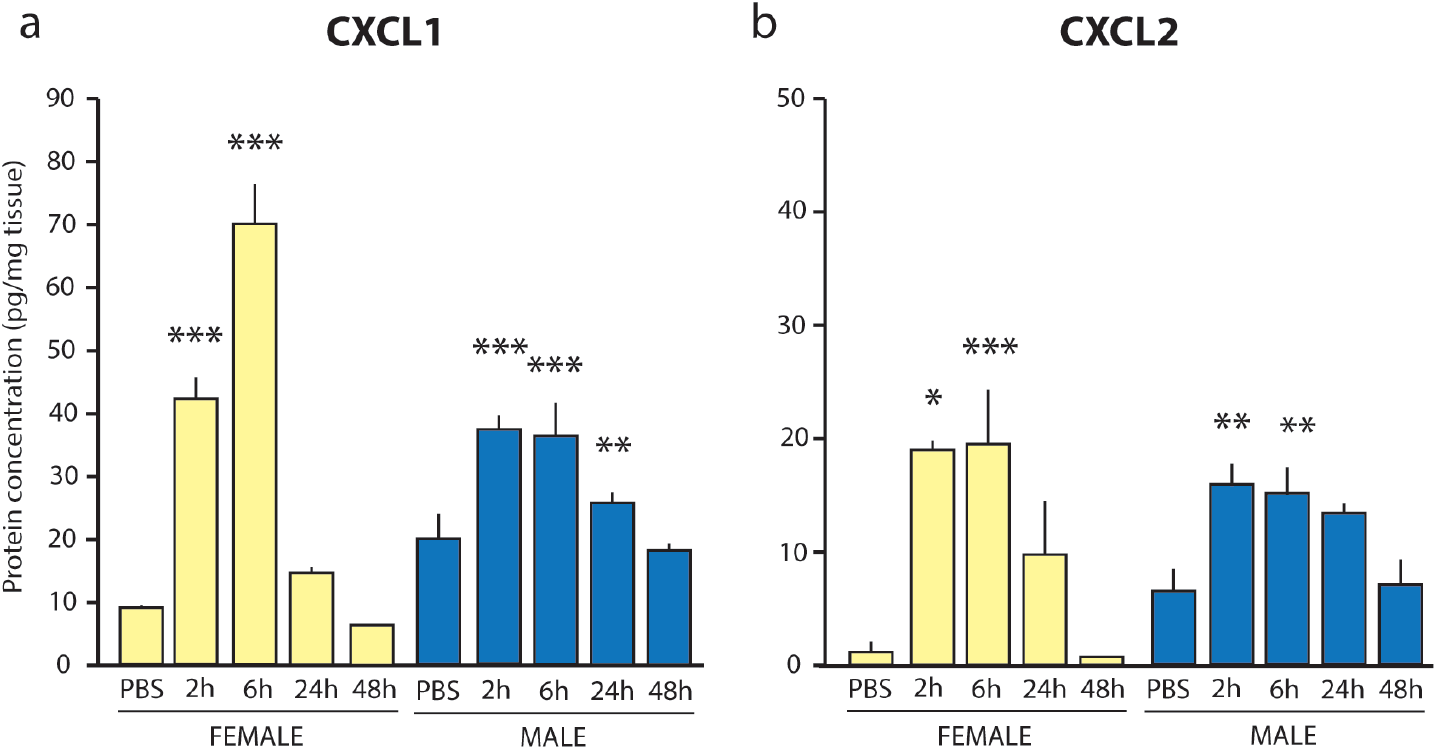
The CXCR2 ligands CXCL1 and CXCL2 (Gro Family). (a) CXCL1 is increased in both males and females 2-6 hours after LPS. In females, this elevation is significantly stronger 6 hours post LPS compared with males. (b) CXCL2 is similarly elevated in both males and females 2-6 hours after LPS. \**p* < 0.05, \*\**p* < 0.01, *** *p* < 0.001 all *cf* PBS.

##### 3.1.2.7 CCR5 Ligands (CCL3, CCL4, CCL5)

The C-C motif chemokines CCL3 and CCL5 also showed sex differences in activity [SEX x TIME: *F*(5,66) = 19.01, *p* < 0.001]. CCL3 and CCL5 were only increased in females with a small but significant increase in CCL3 2 hours after LPS (means ± SEM: PBS: 17.07 ± 1.73 vs 2h: 24.2 ± 1.48; *p* < 0.05), and a 2-fold increase in CCL5 6 hours after LPS (means ± SEM: PBS: 2.35 ± 0.13 vs 6h: 4.38 ± 0.49; *p* < 0.001). CCL4 was not differentially elevated in either sex (Table 1).

#### 3.1.3 Hippocampal cytokine networks

Cytokine networks varied greatly as a factor of sex and time. At rest, male and female cytokine networks were stable to a significance threshold of p=0.01 (Fig. 7a,b). Male networks were characterized by 4 individual clusters of linked cytokines and chemokines with the largest central nodes of this network including CCL5, IL-2, IL-9, and LIF (Fig. 7a). In contrast, the network in females was one large cluster of interconnected cytokines with IL-12p40 serving as the largest central node of this network (Fig. 7b). CSF1 and IL-4 were also highly linked in the female network at rest (Fig. 7b). While the vast majority of cytokine links differed across sexes, links between IL-1α – CCL3 and IL-2 – IL-12p40 were conserved across males and females.

**Figure 7.**
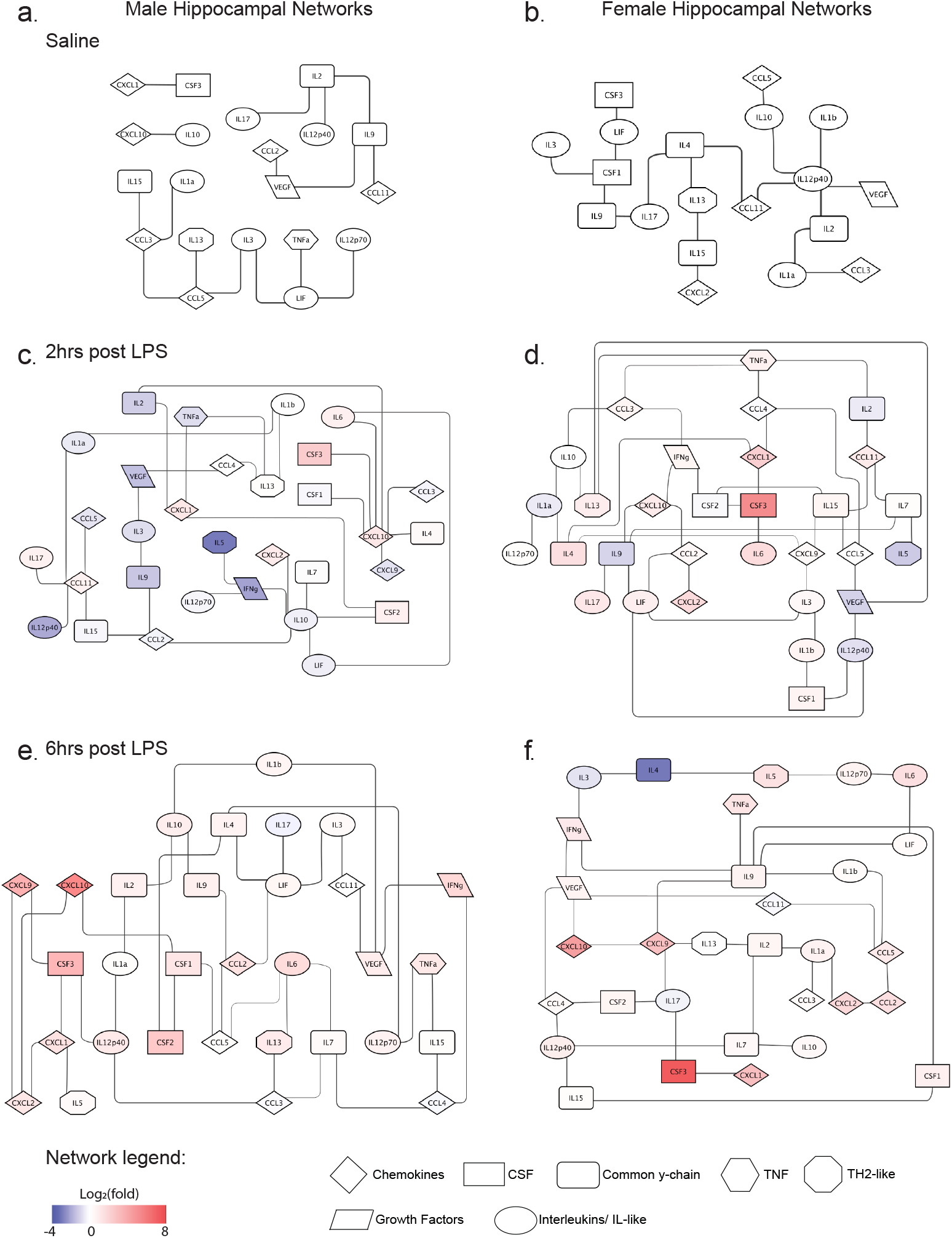
Hippocampal cytokine networks in males and females during LPS-induced cytokine induction. Networks in males (a) and females (b) in saline treated controls. Networks in males (c) and females (d) 2 hours after LPS injection. Networks (e) and females 6 hours after LPS injection. Shape of cytokine nodes correspond to cytokine family and coloration represents relative shifts in concentration (log_2_(fold)) from saline controls.

Cytokine networks increased in complexity and strength starting 2 hours and lasting to 168 hours after injection. At the two-hour time point, CXCL10 was the largest central node of the male network, linking with 7 other cytokines and chemokines (Fig. 7c; significance threshold p=0.0001). CCL11 and IL-10 were the next largest nodes in the network, linking with 4 other cytokines. Similar to raw cytokine concentrations (above), network analysis showed that while some cytokines were increased from saline controls, a large proportion of cytokines were also decreased at this time in males.

However, direct comparisons of log_2_(fold) across all cytokines showed that these changes 2 hours after LPS in males did not significantly differ from saline controls (Fig. 8a; Wilcoxon signed rank test: mean rank=-0.292, confidence interval [CI]=97.06, p=0.281). Unlike the male network, cytokine links were more interconnected in the female network 2 hours post LPS such that TNFα, IL-15, and IL-9, which linked with no more than 4 other cytokines, served as the central nodes of this network (Fig. 7d; significance threshold p=0.0001). Similar to males, several cytokines were reduced compared with saline controls in females 2 hours after LPS injection. However, a larger proportion of cytokines were increased, resulting in a global increase in cytokine concentrations from saline controls (Fig. 8a; Wilcoxon signed rank test: mean rank=0.5242, CI=97.06, p<0.05). Notably, these cytokine levels well exceeded those seen in males (Fig. 8a; two-way ANOVA sex x time interaction: F(4, 292) = 2.809, p<0.05; Bonferroni’s post-test p<0.05). Interestingly, several cytokines were similarly down regulated across sexes at the two-hour timepoint. For example, IL-9, IL-5, VEGF, IL-1α, and IL-12p40 were reduced in both males and females. In addition to conserved effects on individual concentrations, cytokine links between TNFα – IL-13 and IL-15 – CCL11 were also conserved.

**Figure 8.**
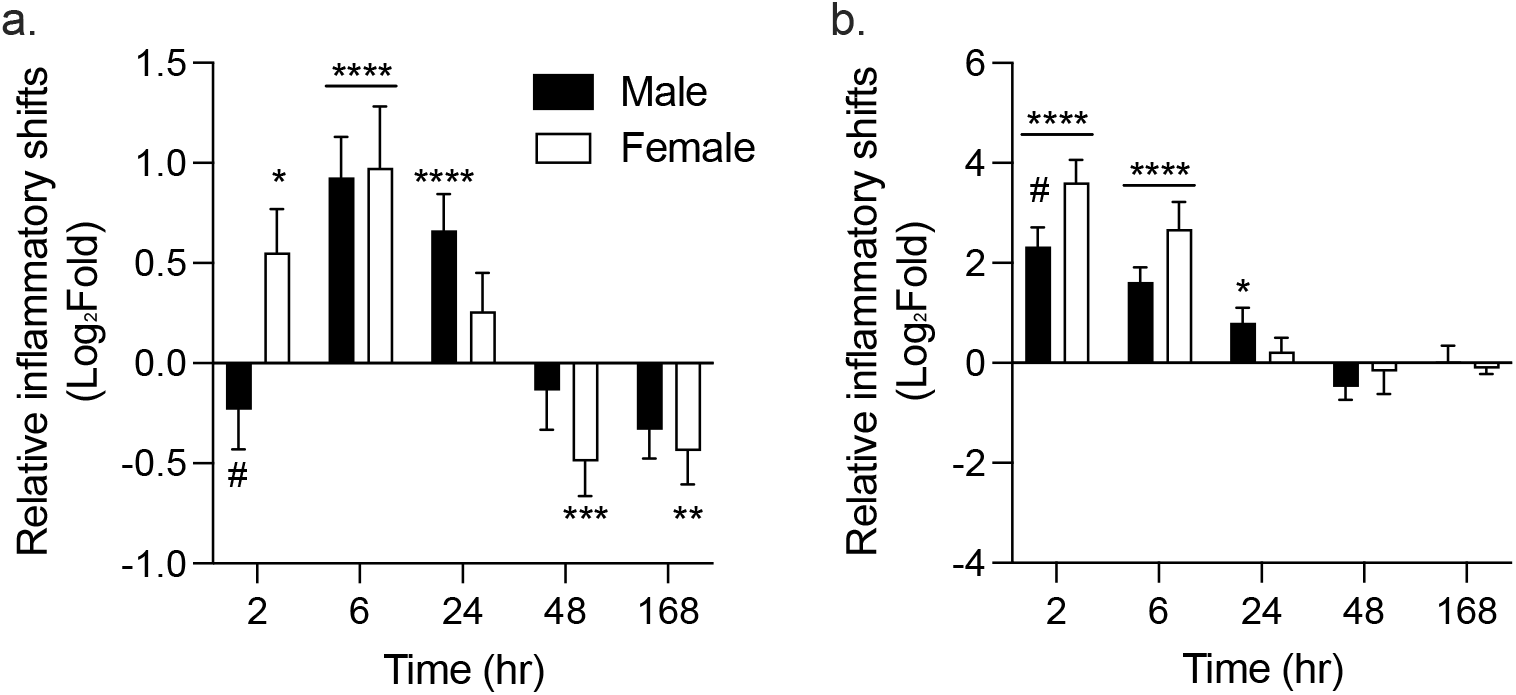
Global shifts in cytokine concentrations 2, 6, 24, 48, and 169 hours after LPS injection relative to saline controls. Relative increases and decreases in cytokine concentrations in the (a) hippocampus and (b) serum. Bonferroni’s post-test: ^#^p<0.05 males vs. females; Wilcoxon signed rank test: *p<0.05, **p<0.01, ***p<0.001, ****p<0.0001 vs. saline controls (0).

By 6 hours after LPS injection, networks in males (Fig. 7e; significance threshold p=0.0001) and females (Fig. 7f; significance threshold p=0.0001) began to look structurally more similar. For example, a larger number of individual cytokine links were conserved across sexes: VEGF – CCL11, CCL2 – CCL5, CXCL1 – CSF3, and IL-2 – IL-1α. Similarly, both networks were characterized by the presence of several smaller interconnected central nodes including CCL5, IL-12p40, and VEGF. Additional central nodes in males 6 hours after LPS included CCL2, CSF3, CXCL2, IL-4, and IL-10. In females, additional central nodes included CXCL9, IL-1α, IL-7, IL-9, IL-17, and INFγ. Outside of structural similarities, global shifts in cytokine concentrations were similarly elevated across males and females at the 6 hours timepoint (Fig. 8a; Bonferroni’s post-test p>0.999) and were significantly elevated from saline controls (Fig. 8a; Wilcoxon signed rank test: males mean rank=0.6468, CI=97.06, p<0.0001; females mean rank=0.6548, CI=97.06, p<0.0001). While global shifts in concentrations were similar, effects on individual cytokines in females were largely varied with some showing increases (CSF3 and CXCL10) and others showing reductions (IL-3 and IL-4) from saline controls. Cytokines in males did not show this same pattern with most cytokines either showing increases or no change from saline controls.

Several similarities in network architecture were also found between males (Fig. 9a; significance threshold p=0.0001) and females (Fig. 9b; significance threshold p=0.0001) 24 hours after LPS injection. For example, links between IL-3 – IL-15, IL-3 – IL-7, IL-12p40 – IL-9, IL-7 – IL-6, CSF2 – CCL4, and CXCL9 – CXCL1 were conserved across networks in males and females. Similarly, IL-3 and CCL2 were centralized nodes in male and female networks at this time. Despite these similarities, there were also striking sex differences. Specifically, global cytokine concentrations in males were still significantly elevated above saline controls (Fig. 8a; Wilcoxon signed rank test: mean rank=0.529, CI=97.06, p<0.0001) while concentrations in females trended toward saline controls (9a; Wilcoxon signed rank test: mean rank=0.245, CI=95.72, p=0.088). These effects in females are likely driven by the strong downregulation of cytokines including IL-3, the single largest central node of the female network at this timepoint (Fig. 9b).

**Figure 9.**
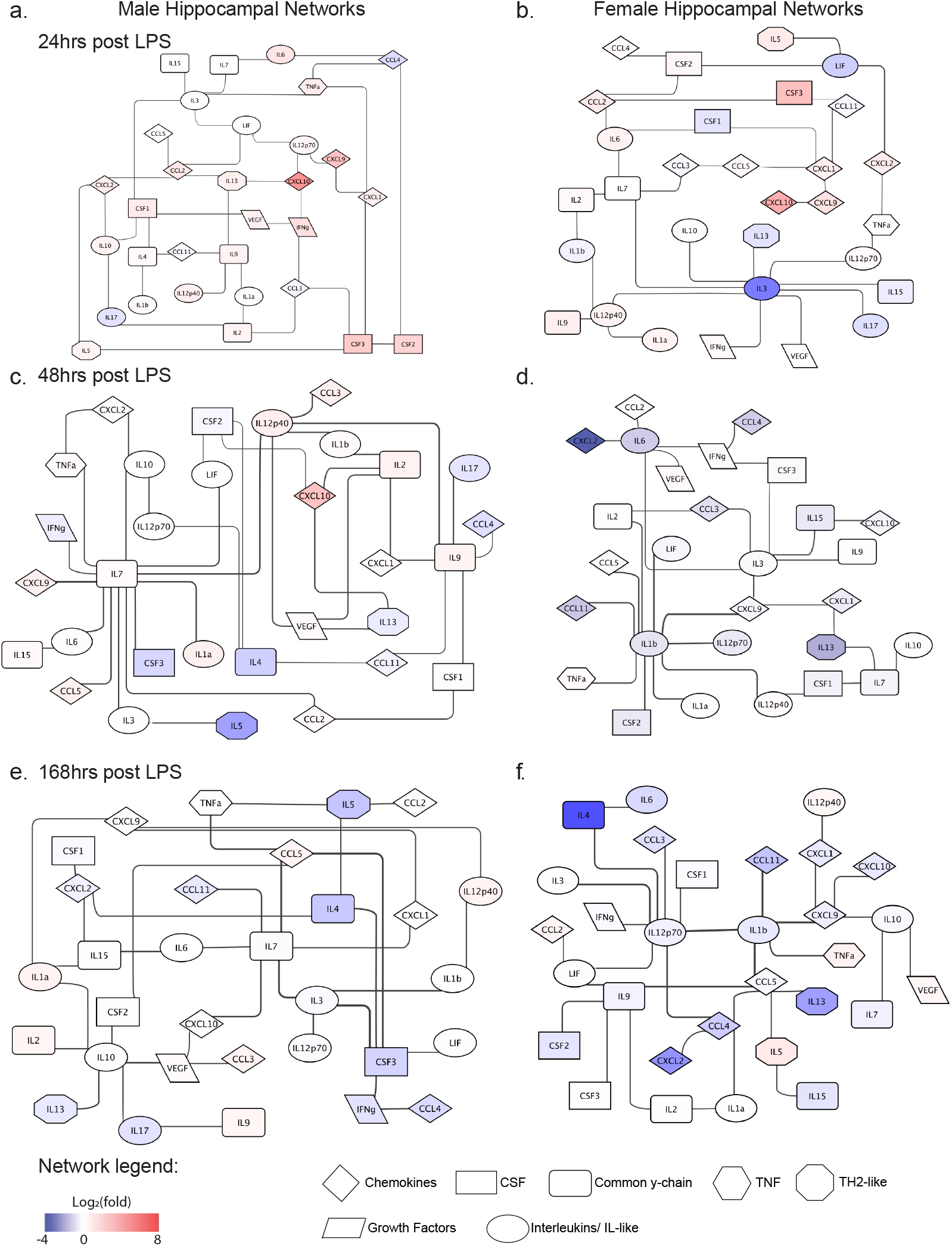
Hippocampal cytokine networks in males and females during the resolution of inflammation. Networks in males (a) and females (b) 24 hours after LPS injection. Networks in males (c) and females (d) 48 hours after LPS injection. Networks (e) and females (f) 168 hours after LPS injection. Shape of cytokine nodes correspond to cytokine family and coloration represents relative shifts in concentration (log_2_(fold)) from saline controls.

In agreement with resolution of inflammation, global cytokine concentrations declined 48 hours after LPS injection. At this timepoint, global concentrations in males were similar to saline controls (Fig. 8a; Wilcoxon signed rank test: mean rank=0.111, CI=97.06, p=0.839). However, global concentrations in females dropped well below the level of saline controls (Fig. 8a; Wilcoxon signed rank test: mean rank: -0.297, CI=95.67, p<0.001). In addition to differences in global concentrations, network architecture began to diverge again in males (Fig. 9c; significance threshold p=0.0001) and females (Fig. 9d; significance threshold p=0.0001). Unlike the 24-hour timepoint, only 3 individual cytokine links were conserved across sexes: IL-1β – IL-12p40, IL-1β – IL-2, and IL-7 – IL-13. Additionally, networks in males primarily centered around IL-7, which was linked with 11 other cytokines, while networks in females centered around IL-1β and to a lesser extent IL-3 and IL-6.

Surprisingly, noted reductions in global cytokine concentrations in females were still detected 168 hours after the LPS injection (Fig. 8a; Wilcoxon signed rank test: mean rank=-0.288, CI=95.72, p<0.01). In contrast, while several individual cytokines were still reduced in males, these effects did not result in significant reductions of global cytokine concentrations compared with saline controls (Fig. 8a; Wilcoxon signed rank test: mean rank=-0.0371, CI=97.06, p=0.182). Similar to the 48-hour timepoint, only three cytokine links were conserved across male (Fig. 9e; significance threshold p=0.0001) and female (Fig. 9f; significance threshold p=0.0001) networks: IL-12p70 – IL-3; CXCL9 – CXCL1; and IL-10 – VEGF. Further, male networks were strongly centered around CSF3, IL-10, IL-7, and CCL5 (Fig. 9e). While female networks were centered around IL-12p70, IL-1β, IL-9, and CCL5 (Fig. 9f).

### 3.2 Peripheral cytokines

Of the 32 cytokines tested, 26 exhibited a significant elevation in the periphery after systemic LPS injection (Main effect of TIME, Table 1), six (IL-2, IL-9, CSF1, CSF3, CXCL5, and VEGF) were not significantly elevated when collapsed across males and females (Table 2). Compared with hippocampal cytokines, activation of peripheral cytokines was more robust, with stronger effect sizes and more rapid induction and resolution after LPS. Post-hoc tests showed that independent of sex, all but one activated cytokine was upregulated 2 hours after LPS (all *p* < 0.001), with CXCL9 upregulated only after 6 hours (2h: *p* = 0.696; 6h: *p* < 0.001). At 6 hours post LPS, CSF2 (*p* < 0.001), TNFα (*p* < 0.05), IL-1β (*p* < 0.001), IL-10 (*p* < 0.001), IL-12p40 (*p* < 0.001), CXCL10 (*p* < 0.001), CCL2 (*p* < 0.001), CCL3 (*p* < 0.001), and CCL5 (*p* < 0.001) remained elevated. With the exception of CXCL9 (24h: *p* < 0.001) and CSF2 (24h: *p* < 0.05) all cytokines had returned to baseline by 24 hours post LPS injection.

**Table 2.** Serum cytokines: Statistical analysis Summary. Overall multivariate ANOVA results for the Main effect of TIME and the interaction effect (SEX x TIME).\**p* < 0.05, \*\**p* < 0.01, \*\*\**p* < 0.001.

| CYTOKINE | MAIN EFFECT: TIME | INTERACTION: SEX x TIME |
| --- | --- | --- |
| **Serum cytokines differentially activated in males and females** |  |  |
| **IL-1β** | *** $F_{(5,66)} = 88.05$ $\eta^2_p = 0.87$ | ** $F_{(5,66)} = 4.25$ $\eta^2_p = 0.24$ |
| **CCL3 (MIP1α)** | *** $F_{(5,66)} = 124.39$ $\eta^2_p = 0.90$ | ** $F_{(5,66)} = 3.39$ $\eta^2_p = 0.20$ |
| **CCL4 (MIP1β)** | *** $F_{(5,66)} = 25.10$ $\eta^2_p = 0.66$ | *** $F_{(5,66)} = 5.67$ $\eta^2_p = 0.30$ |
| **CCL5 (RANTES)** | *** $F_{(5,66)} = 32.56$ $\eta^2_p = 0.71$ | *** $F_{(5,66)} = 10.08$ $\eta^2_p = 0.50$ |
| **CCL11 (Eotaxin)** | ** $F_{(5,66)} = 4.78$ $\eta^2_p = 0.27$ | * $F_{(5,66)} = 2.87$ $\eta^2_p = 0.18$ |
| **CXCL2 (MIP2)** | *** $F_{(5,66)} = 214.58$ $\eta^2_p = 0.94$ | ** $F_{(5,66)} = 3.70$ $\eta^2_p = 0.22$ |
| **CXCL10 (IP-10)** | *** $F_{(5,66)} = 64.82$ $\eta^2_p = 0.83$ | ** $F_{(5,66)} = 4.27$ $\eta^2_p = 0.24$ |
| **Serum cytokines activated in both sexes** |  |  |
| **IL-1α** | * $F_{(5,66)} = 2.53$ $\eta^2_p = 0.16$ | n.s. |
| **TNFα** | *** $F_{(5,66)} = 230.87$ $\eta^2_p = 0.95$ | n.s. |
| **IL-6** | *** $F_{(5,66)} = 338.02$ $\eta^2_p = 0.96$ | n.s. |
| **IL-10** | *** $F_{(5,66)} = 71.24$ $\eta^2_p = 0.84$ | n.s. |
| **IL-4** | *** $F_{(5,66)} = 6.39$ $\eta^2_p = 0.33$ | n.s. |
| **IL-13** | *** $F_{(5,66)} = 23.28$ $\eta^2_p = 0.64$ | n.s. |
| **IL-15** | *** $F_{(5,66)} = 16.26$ $\eta^2_p = 0.55$ | n.s. |
| **IFNγ** | * $F_{(5,66)} = 2.53$ $\eta^2_p = 0.16$ | n.s. |
| **CXCL9 (MIG)** | *** $F_{(5,66)} = 69.68$ $\eta^2_p = 0.84$ | n.s. |
| **CSF2 (GM-CSF)** | *** $F_{(5,66)} = 50.16$ $\eta^2_p = 0.79$ | n.s. |
| **CXCL1 (KC)** | *** $F_{(5,66)} = 121.07$ $\eta^2_p = 0.90$ | n.s. |
| **CCL2 (MCP1)** | *** $F_{(5,66)} = 59.97$ $\eta^2_p = 0.82$ | n.s. |
| **IL-3** | *** $F_{(5,66)} = 6.63$ $\eta^2_p = 0.33$ | n.s. |
| **IL-5** | *** $F_{(5,66)} = 5.77$ $\eta^2_p = 0.30$ | n.s. |
| **IL-7** | *** $F_{(5,66)} = 21.25$ $\eta^2_p = 0.62$ | n.s. |
| **IL-12p40** | *** $F_{(5,66)} = 63.82$ $\eta^2_p = 0.83$ | n.s. |
| **IL-12p70** | *** $F_{(5,66)} = 12.52$ $\eta^2_p = 0.49$ | n.s. |
| **IL-17** | *** $F_{(5,66)} = 18.09$ $\eta^2_p = 0.58$ | n.s. |
| **LIF** | *** $F_{(5,66)} = 66.31$ $\eta^2_p = 0.83$ | n.s. |

#### 3.2.1 Sex differences in activation of serum cytokines

In the serum, 7 of 32 cytokines, IL-1β, CCL3, CCL4, CCL5, CCL11, CXCL2, and CXCL10, showed differential activation in males and females, with a significant SEX x TIME interaction (see Table 2). CCL11 showed very small effect size for SEX x TIME interaction (η^2^p = 0.18) and was not further examined. Unlike hippocampal cytokines that showed sex differences in magnitude, time course, and pattern of cytokines, serum cytokines only showed sex differences in magnitude of activation. Furthermore, relatively few cytokines in serum compared with hippocampus showed sex differences in activation after LPS injection.

IL-1β was significantly elevated at 2 and 6 hours after LPS injection in both sexes [TIME: *F*(5,66) = 88.05, *p* < 0.001; SEX x TIME: *F*(5,66) = 4.25, *p* < 0.001; 2h: all *p* < 0.001; 6h: all *p* < 0.001), with significantly greater levels in females at 6 hours compared with males (*p* < 0.05) (Fig. 10a). In contrast, TNFα was elevated 2 and 6 hours after LPS [TIME: *F*(5,66) = 230.87, *p* < 0.001; 2h: *p* < 0.001; 6h: *p* = 0.018] (Fig. 10b) and IL-6 at 2 hours after LPS injection [TIME: *F*(5,66) = 230.87, *p* < 0.001; 2h: *p* < 0.001] with no differences between males and females [SEX x TIME: TNFα: *F*(5,66) < 1; IL-6:*F*(5,66) = 1.33; *p* = 0.09] (Fig. 10c).

**Figure 10.**
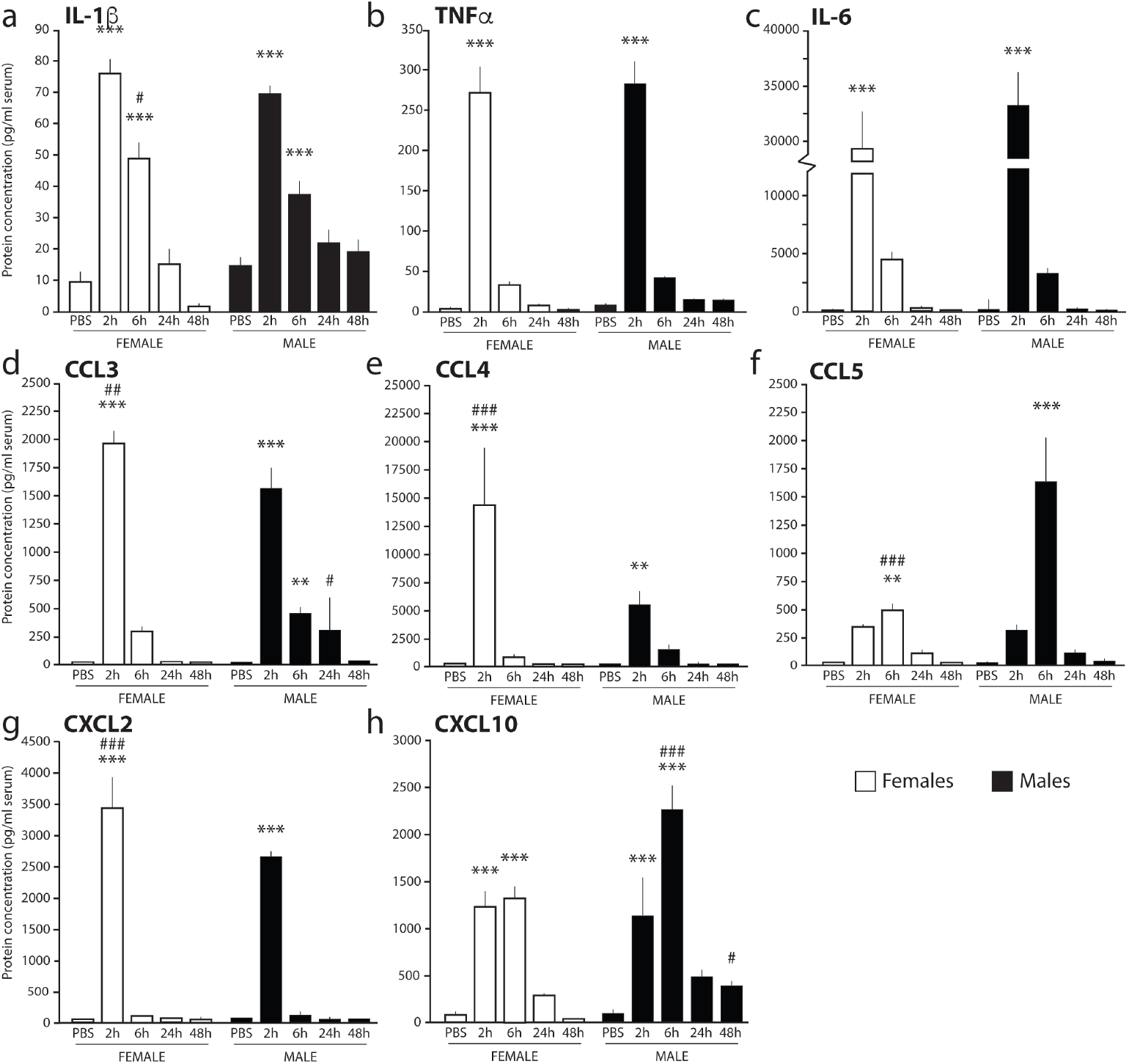
Serum cytokines in males and females after systemic LPS injection. (a) IL-1β is increased 2 and 6 hours after LPS in females and males. (b) TNFα is elevated 2 hours after LPS in both sexes. (c) IL-6 is increased 2 hours after LPS in males and in females. (d) CXCL2 is increased 2 hours after LPS in both sexes, and more strongly in females. (e) CXCL10 is increased 2-6 hours after LPS in both sexes, and more strongly in males. (f) CCL3 is elevted more strongly in females 2 hours after LPS, and more persistently in males. (g) CCL4 is more strongly elevated in females 2 hours after LPS. (h) CCL5 is elevated more strongly in males 2 hours after LPS. \**p* < 0.05, \*\**p* < 0.01, \*\*\**p* < 0.001 all cf PBS; #*p* < 0.05, ### *p* < 0.001, male vs female.

CXCL2 was elevated in both sexes at 2 hours [TIME: *F*(5,66) = 213.58; *p* < 0.001; SEX x TIME: *F*(5,66) = 3.70, *p* = 0.005; 2h: all *p* < 0.001), with stronger activation in females compared with males (*p* < 0.001) (Fig. 10d). In contrast, CXCL10 was elevated in both sexes at 2 and 6 hours [TIME: *F*(5,66) = 64.82; *p* < 0.001; SEX x TIME: *F*(5,66) = 4.27, *p* = 0.002; 2h: all *p* < 0.001; 6h: all *p* < 0.001], with stronger activation in males than females 6 hours after LPS injection (*p* < 0.001) (Fig. 10e).

The most striking sex differences were observed in the CCR5 ligands, CCL3, CCL4 and CCL5. Both males and females showed activation of CCL3 in serum 2 hours after LPS [TIME: *F*(5,66) = 124.39; *p* < 0.001; SEX x TIME: *F*(5,66) = 3.39, *p* = 0.009; 2h: male: *p* < 0.001; female *p* < 0.001] with stronger levels of CCL3 in females (*p* < 0.01). In contrast, only males showed activation 6 hours after LPS (6h: male *p* < 0.01; female *p* = 0.21), and this activation remained higher than that of females 24 hours after LPS (*p* < 0.05) (Fig. 10f). In addition, CCL4 [TIME: *F*(5,66) = 25.10; *p* < 0.001; SEX x TIME: *F*(5,66) = 5.67, *p* = 0.009; 2h: male: *p* < 0.01; female *p* < 0.001] and CCL5 [TIME: *F*(5,66) = 32.56; *p* < 0.001; SEX x TIME: *F*(5,66) = 10.08, *p* < 0.001; 6h: male: *p* < 0.001; female *p* = 0.007] showed opposing patterns, with higher CCL4 levels in females compared with males 2 hours after LPS (*p* < 0.001) (Fig. 10g); and lower levels of CCL5 6 hours after injection (*p* < 0.001) (Fig. 10h).

#### 3.2.2 Peripheral cytokine networks

Serum cytokine concentrations were also used for network analysis. At rest (saline controls), cytokine networks in males (Fig. 11a) and females (Fig. 11b) were stable to a significance threshold of p=0.01 and were characterized by individual clusters of linked cytokines. The largest network cluster in males was characterized by the chemokines CXCL9, CCL11, CCL2, CCL3, and CCL5 that converged on CXCL10. A second smaller cluster was also detected in which IL-13 and IL-6 converged onto CXCL1. Network analysis in females also found three individual clusters of linked cytokines. The largest of these clusters was primarily comprised of the interleukins IL-4, IL-10, and IL-1β and LIF which converged onto IL-12p70. Notably, links between CXCL10 and CCL11 were conserved across sexes and in females was the single smallest cluster at this time point.

**Figure 11.**
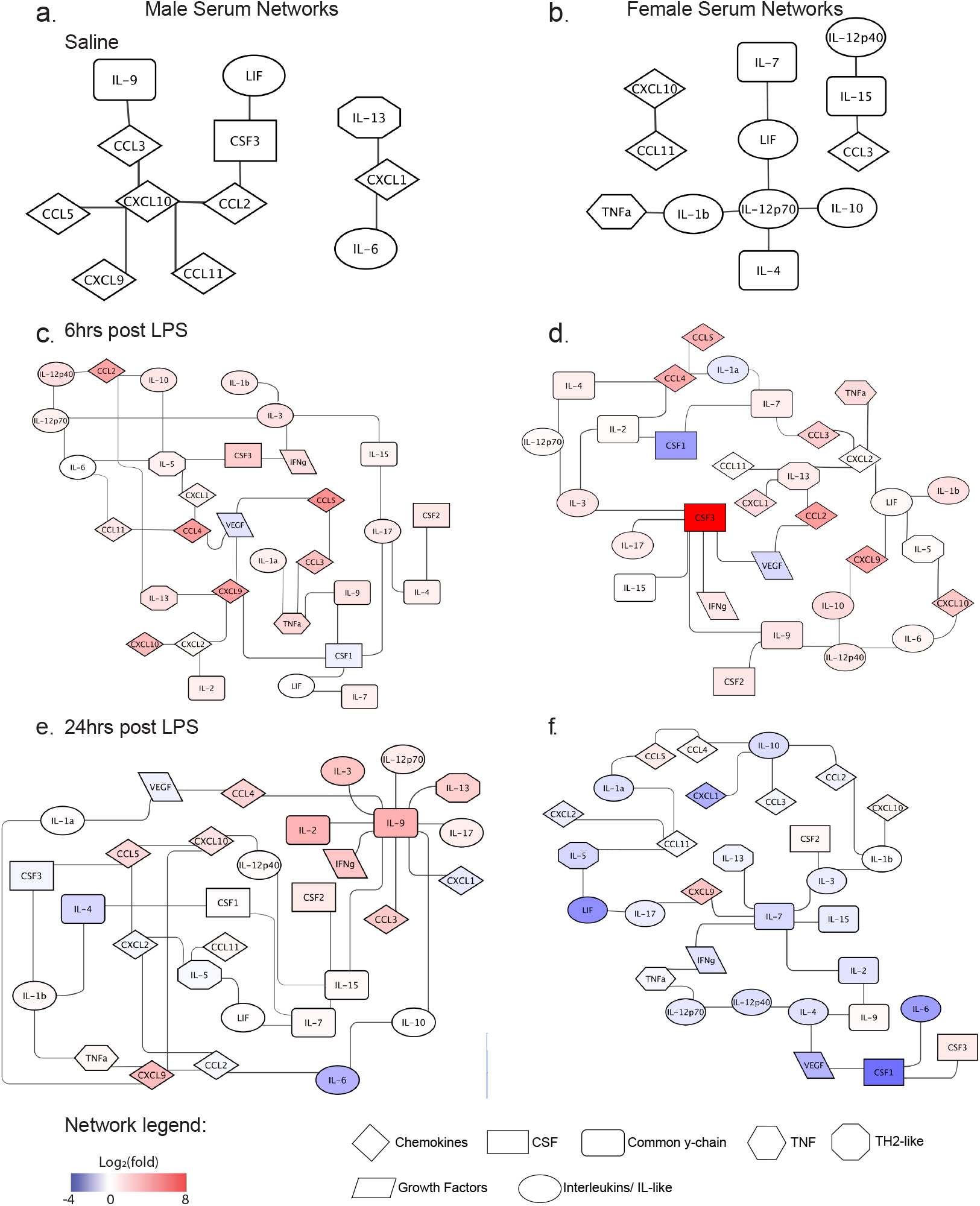
Serum cytokine networks in males and females during LPS-induced cytokine induction. Networks in males (a) and females (b) in saline treated controls. Networks in males (c) and females (d) 6 hours after LPS injection. Networks (e) and females (f) 24 hours after LPS injection. Shape of cytokine nodes correspond to cytokine family and coloration represents relative shifts in concentration (log_2_(fold)) from saline controls

No networks were detected at the 2-hour time point in either males or females (significance thresholds tested p ≤ 0.07). While no networks were detected, global cytokine concentrations were elevated in both males (Fig. 8b; Wilcoxon signed rank test: mean rank = 2.0, CI = 98.0, p<0.0001) and females (Fig. 8b; Wilcoxon signed rank test: mean rank = 2.9, CI – 98.0, p<0.0001) above saline controls. Additionally, global shifts in cytokine concentrations were greater in females vs. males (Fig. 8b; two-way ANOVA sex x time interaction: F(4, 308) = 2.48, p<0.05; Bonferroni’s post-test p=0.05), supporting a role for enhanced inflammatory reactivity in females.

Networks were once again detected at 6 hours in both males (Fig. 11c) and females (Fig. 11d), which were both stable to a significance threshold of p=0.0001. Unlike networks at rest, networks 6 hours after LPS injection were characterized by several interconnected central nodes. In males, the largest of these central nodes included CSF1, CXCL9, IL-3, and IL-5 (Fig. 11c). Networks in females at this time were also characterized by several interconnecting central nodes (Fig. 11d). The largest central node of the female network at this time was CSF3 which was directly linked with IL-3, IL-9, IL-15, IL-17, INFγ, and VEGF. Interestingly, cytokine networks were most different between males and females at this time point with only one cytokine link conserved across sexes: CSF3-INFγ. In addition to noted differences in cytokine connectivity, females demonstrated the broadest range of cytokine concentrations compared with males: reaching a log_2_(fold) of 10.8 for CSF3 and -4.1 for CSF1 in females versus 5.8 for CCL5 and -1.2 for VEGF in males. Despite differences in the overall ranges of individual cytokine concentrations, global cytokine concentrations did not differ between males and females 6 hours after LPS injection (Fig. 8b, Bonferroni’s post-test p=0.17).

Similar to the 6-hour time point, networks in males (Fig. 11e) and females (Fig. 11f) 24 hours after LPS injection were characterized by a series of interconnected central nodes. Both networks were stable down to a significance threshold of p=0.0001. In males, IL-9 was the single largest central node of the network and was also one of the most up-regulated cytokines at this time point (log_2_(fold) = 4.2). Smaller central nodes of this network included pro-inflammatory IL-15 and CXCL9 which were respectively linked to CSF2, IL-9, IL-7, IL-12p40 and IL-1α, CXCL10, TNFα, CCL2. While anti-inflammatory IL-9 was the largest central node in the male network 24 hours after LPS, pro-inflammatory IL-7 was the largest central node in the female network at this time point (Fig. 11f). In females IL-7 was directly linked with 6 other cytokines: CXCL9, IL2, IL-3, IL-13, IL-15, and INFγ. IL-10 was the next largest node of the network with four connections to CCL2, CCL3, CCL4, and CXCL1. Unlike networks at 6 hours, several cytokine links were conserved across male and female networks 24 hours after LPS injection: IL-15 – IL-7, IL-5 – CCL11, IL-5 – LIF, IL-9 – IL-2. Similar to effects in the hippocampus, males showed sustained increases in global cytokine concentrations 24 hours after LPS injection (Fig. 8b, Wilcoxon signed rank test: mean rank = 0.4, CI = 98.0, p<0.05), while females no longer showed overall increases in cytokine levels as compared to saline controls (Fig. 8b; Wilcoxon signed rank test: mean rank = 0.16, CI = 98.0, p=0.54). However, no statistical differences were detected in overall cytokine concentrations between males and females (Fig. 8b; Bonferroni’s post-test: p=0.99).

In agreement with resolution of inflammation, global cytokine concentrations in both males (Fig. 8b; Wilcoxon signed rank test: mean rank = 0.06, CI = 98.0, p = 0.39) and females (Fig. 8b; Wilcoxon signed rank test: mean rank = -0.14, CI = 95.7, p = 0.52) were comparable to saline controls 48 hours after LPS injection. At this time, male and female cytokine networks were stable to a significance threshold of p = 0.0001 (Fig.13a, b). Networks in males were quite sparse with central nodes sharing no more than 4 connections with other cytokines and chemokines (Fig. 12a). These central nodes included IL-15 and IL-17, which were respectively linked to CCL3, CXCL2, IL-10, IL-12p40 and CCL4, CCL5, IL-1β, and IL-4 (Fig. 12a). Unlike networks in males where central nodes only had sparse connectivity, networks in females largely centered around CSF1 which linked with 9 additional cytokines: CCL3, CXCL1, IL-1α, IL-1β, IL-2, IL-7, IL-10, IL-12p40, and INFγ (Fig. 12b). Given the vast differences in network architecture across sexes, it was not surprising that only 2 cytokine links were conserved across sexes: CCL4 – VEGF and CSF1 – IL-7.

**Figure 12.**
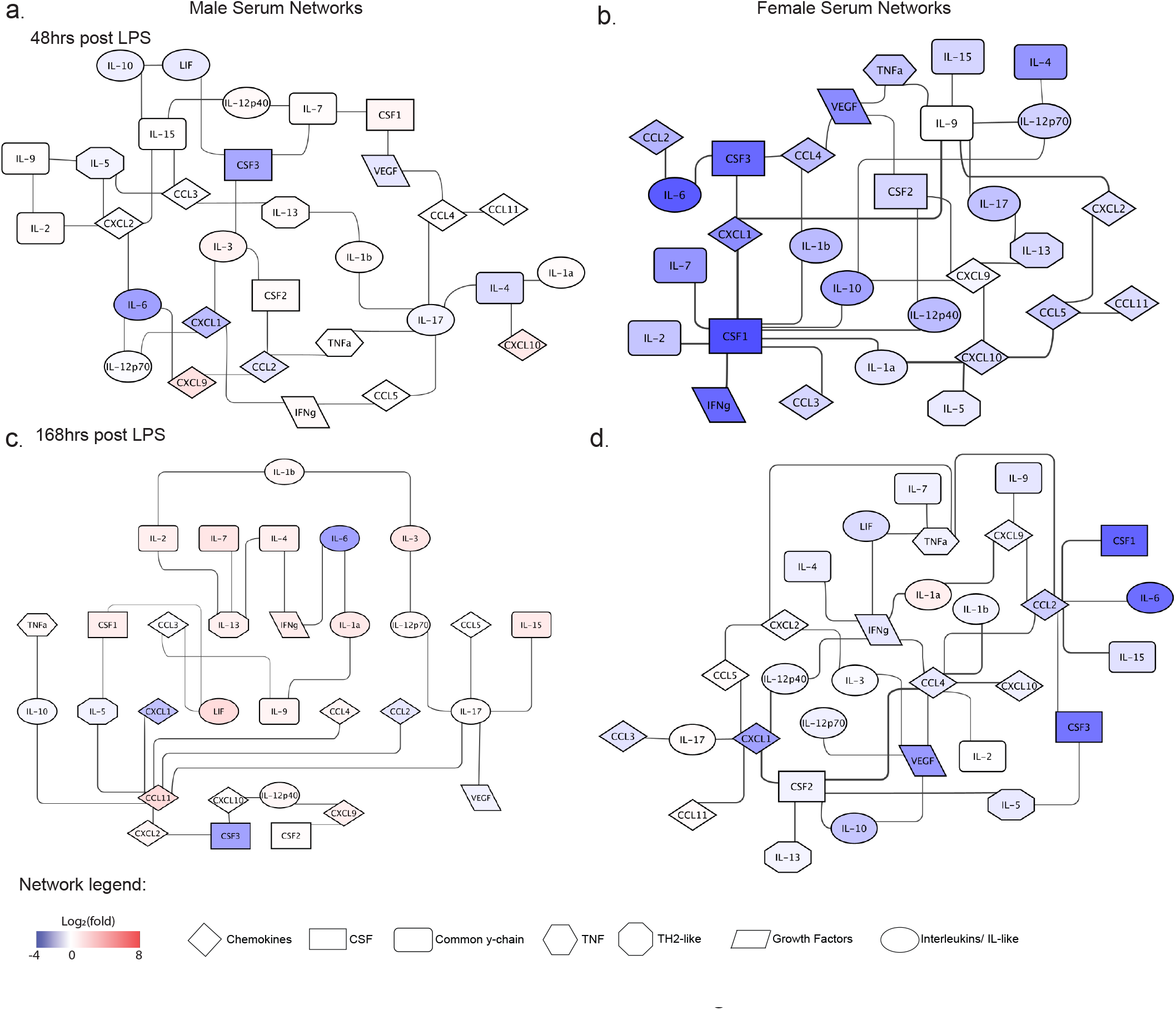
Serum cytokine networks in males and females during the resolution of inflammation. Networks in males (a) and females (b) 48 hours after LPS injection. Networks in males (c) and females (d) 168 hours after LPS injection. Shape of cytokine nodes correspond to cytokine family and coloration represents relative shifts in concentration (log_2_(fold)) from saline controls.

Similar to the 48-hour time point, global cytokine concentrations in the serum were unchanged from saline controls 168 hours after LPS injection in in both males (Fig. 8b; Wilcoxon signed rank test: mean rank = 0.47, CI = 98.0, p = 0.07), and females (Fig. 8b; Wilcoxon signed rank test: mean rank = -0.12, CI = 98.0, p=0.38). Networks in males and females were still stable to a significance threshold of p=0.0001, and were largely characterized by interconnected central nodes (Fig. 12c,d). In males, the single largest central node was CCL11 which was linked with CCL2, CCL4, CSF3, CXCL1, CXCL2, IL-5, IL-10, and IL-17 (Fig. 12c). In contrast, networks in females largely centered around CXCL1, CCL2, CCL4, and CSF2 (Fig. 12d). The vast majority of these central nodes in female networks had 5-6 cytokine links which included a direct link to one or more central nodes of this network. Despite noted differences in network architecture, links between IL-4 – INFγ and CXCL1 – CCL11 were conserved across sexes. Interestingly, links between CXCL1 with CCL11 were also conserved across sexes in the resting saline condition and may signal that responses to LPS have completely resolved.

## 4. DISCUSSION

This study demonstrated sex differences in hippocampal cytokine networks activated after a systemic LPS challenge. We observed differences across production, magnitude, and rate of resolution of neuroimmune levels across the one-week time course. For example, male-specific activation was observed for CSF1 and CSF2, IFNɣ, and IL-10, with stronger activation in CXCL9 and 10. In contrast, female-specific activation was observed for IL-2 and IL-15, CCL3 and 5, IL-1α and IL-4, with stronger activation of CSF3 and CXCL1. These differences emerged amongst a robust hippocampal cytokine response in both sexes.

The hippocampal cytokine response was distinct from the peripheral immune response, as measured by direct comparisons between cytokine levels and by network analyses. Cytokine activation was slower and more persistent in the hippocampus compared with the periphery; and notably, IL-1β, IL-6, and TNFα were elevated to a lesser degree in the hippocampus than in the periphery. Consistent with other work ^18,54– 56^, hippocampus had proportionally greater activation of cytokines including CXCL9, CXCL10, and CSF3. The fundamental differences in cytokine network architecture between serum and hippocampus strongly suggests that cytokine activity in the brain is not simply a recapitulation (or artefact) of circulating cytokine levels, but rather represents central immune activation and regulation. Unlike hippocampus, in which we observed sex differences in time course, pattern and magnitude, serum cytokines of males and females differed primarily in magnitude. Both sexes showed elevated levels of the same cytokines, but females had stronger activation of IL-1β, CXCL2, CCL3 and CCL4, whereas males showed stronger activation of CCL5 and CXCL10. As expected^57– 59^, this was also accompanied by robust sex differences in cytokine network architecture across all timepoints.

Males and females also exhibited distinct kinetics of hippocampal cytokine regulation after systemic LPS. Females showed a rapid elevation of cytokines that peaked at 2-6 hours and trended toward resting baseline levels at 24 hours. In contrast, males showed slower activation, with all cytokines elevated at 6 hours and most persisting for at least 24 hours. Faster neuroimmune activation and resolution is consistent with more efficient immune response by females compared with males in response to toll like receptor (TLR) activation in the periphery ^60^. Global shifts in cytokine concentrations also demonstrated a period of down regulation of cytokines in the hippocampus 48 hours after LPS injection. While these effects were evident in both sexes, cytokine downregulation was greatest and most sustained in females. This new observation may represent a refractory period and may be a window of immune vulnerability that is greater in females compared with males.

The overall structures of cytokine networks in the hippocampus were statistically robust in both sexes, and differed markedly in links between individual cytokines and chemokines. These differences in network structure influence the functional outcome of immune activation. At rest, networks in males favored CCL3, CCL5, IL-2, IL-9, and LIF. With exception of IL-9 and LIF, the functions of these cytokines are skewed toward the recruitment of peripheral inflammatory cells to the brain^61–64^, disruption of the blood brain barrier (BBB) ^65^, and the activation of T-cells ^66^. In contrast, networks in females at this same time point were centered around CSF1, IL-4, and IL-12p40, which are generally involved in the recruitment of macrophages ^67^, the activation and proliferation of microglia ^68^, and, in the case of IL-4, inhibition of NF-kB ^69^ and microglial polarization toward M2 anti-inflammatory phenotypes ^70^.

These cytokine networks reflect both the individual shifts in cytokine concentrations observed in the current study and are highly consistent with previous work demonstrating the importance of IL-4 in microglia of females after ischemic stroke ^71^. These findings are consistent with a number of previously observed sex differences, including that males have a greater IL-10 response compared with females ^72^ and that IL-2 ^18^ and IL-13 ^19^ play different roles in the hippocampus in males compared with females.

Sex differences in patterns of cytokines have implications for what neuroimmune cells are recruited in males compared with females. For example, sex differences in IL-4, IL-13, and IFNɣ are of particular interest for their differential roles in microglia activation. IFNɣ has been proposed to cause microglia polarization to a M1-like (“inflammatory”) pattern of cytokine response, and IL-4/IL-13 towards M2-like (“alternative”) microglial response ^73^, similar to macrophage M1/M2 states ^74^. The male-specific increase in CSF2 may contribute to the more persistent inflammatory response in males ^75^, and in females, the short-lasting IL-1β, IL-6 and TNFα profile suggests rapid activation of regulatory processes.

Network analyses supported a role for microglia in hippocampal cytokine activation, and are consistent with shifts in M1-/M2-like polarization. In males, for example, hippocampal cytokine networks 2 hours after LPS showed strong links between CCL3, CXCL9, and IL-6 all of which converge onto CXCL10, the largest central node of the network. These cytokines can all be released by M1-type microglia^76^. Thus, strong MI scores between CXCL10, CCL3, CXCL9, and IL-6, suggest co-release of these cytokines and implicate M1-skewed microglia as the dominant cell type contributing to the cytokine response 2 hours after LPS injection in males.

The phenotypic classification of microglia as M1- or M2-is clearly an oversimplification^77^. Instead, recent work has begun to show that microglia exist in a variety of intermediate states that do not adhere to these strict classifications. Specifically, recent work by Hammond et al. revealed at least nine distinct microglial states that were coupled with unique gene expression profiles ^78^, highlighting the complexity and heterogeneity of microglia. Hippocampal cytokine network analyses provided support for such intermediate microglial states in males and females. Networking of IL-12p40 with IL-1β, IL-2, CCL11, IL-10, and VEGF in females under saline control conditions provide the clearest example of this phenomenon. With exception of VEGF, all of these cytokines can be released from microglia. Under the “traditional” phenotypic classifications, M1-type are known to release IL-12p40, IL-1β, and IL-2, while M2-type release CCL11 and IL-10 ^76^. Thus, clustering of and strong MI scores of cytokines linked with IL-12p40 are suggestive of co-release from microglia with intermediate (M1/M2) phenotypes in females.

Cytokine network analyses, using comparisons of overlapping cellular sources of central nodes and directly linked cytokines, suggest that microglia were the primary drivers of early phase neuroimmune response to LPS. However, hippocampal cytokine networks in both sexes became increasingly indicative of T-cell and astrocytic contributions at 6 and 24 hours after LPS injection, indicating a switch from a predominantly microglia-dominated immune response to a T-cell- and astrocyte-dominated response. Similarly, by 48 hours and persisting to 168 hours, cytokine networks were predominantly focused around astrocytic and dendritic cytokines and chemokines with some contribution of microglia and T-cells. This specific temporal pattern of cellular activation described by cytokine networks is consistent with previously published work showing rapid microglial activation ^79,80^, transition toward M2 favoring states at 24 hours^81^, T-cell recruitment to mesenchymal compartments during the early phase of the inflammatory response^82^, and latent astrocyte activation beginning 12 hours following LPS injection ^83^. Together with these previous data, our network analyses provide strong empirical basis for hypothesis-based experiments aimed to directly test these associations between neuroimmune cells, cytokines, and sex.

The implications of these cytokine networks for the involvement of various immune cell types are consistent with recent work on neuroimmune activation. Astrocytes, mast cells, eosinophils, leukocytes and macrophages are actively involved in – and contribute to sex differences in – cytokine production and network dynamics. Astrocytes, for example, show sex-specificity of cytokine release in response to LPS or injury ^84,85^. CXCL1 and CXCL2 play key roles in recruitment of immune cells including leukocytes ^86^ and neutrophils ^87^ during neuroimmune activation. The greater activation of CXCL1 in females compared to males suggests that differential immune cell recruitment and activation may be both cause and consequence of sex differences in cytokine network activation. This network analysis begins to describe precise, sex-specific patterns cytokine release in the brain, and suggests temporal dynamics of microglia, astrocytes and other infiltrating immune cells.

In this project, we demonstrated that a systemic LPS challenge activates broad sex-specific cytokine profiles from multiple families in C57Bl6/N mice in both hippocampus and in the periphery. Importantly, hippocampal cytokine profiles were distinct from those observed from serum, demonstrating that the neuroimmune response is not a direct result of LPS-induced increases in peripheral cytokines. This distinct hippocampal neuroimmune response shows sex differences in the families of cytokines activated, the magnitude of activation, and the time course of activation and regulation of cytokine signaling. Combining comparisons of individual cytokines with network analyses provides a broader context to understand immune effectors across sex. Here, cytokine network analysis suggested that these different patterns of cytokines may mediate differential cell recruitment, microglia polarization, and intracellular signaling, which may lead to very different functional outcomes of signaling in males and females. Delineating network dynamics of cytokine signaling in the hippocampus, and understanding the interactions and relationships between individual cytokines in males and in females will thereby provide a new framework for understanding sex-biased susceptibility to psychiatric and neurological disorders.

## Supporting information

Network Links Data_Supp Tables

## FUNDING AND ACKNOWLEDGEMENTS

This research was funded by R00 MH093459 to NCT and T32-DK101357 to JEF. We would like to acknowledge Drs. Katie Collette, Lacie Turnbull, Ashley Keiser, Daria Tchessalova, and Caitlin Posillico for their helpful comments on an earlier version of this manuscript.

