## Supplementary material for "Sex differences in hippocampal cytokine networks after systemic immune challenge": Network Links Data_Supp Tables

| | Males | | Females | |
|---|---|---|---|---|
| | Cytokine Links | MI Score | Cytokine Links | MI Score |
| Hippocampus | CCL2 – VEGF | 0.5063 | CCL3 – IL-1a * | 0.4867 |
| | CCL3 – CCL5 | 0.5123 | CSF1 – IL-3 | 0.5298 |
| | CCL3 – IL-15 | 0.5444 | CSF1 – IL-9 | 0.4982 |
| | CCL3 – IL-1a * | 0.5808 | CSF1 – LIF | 0.5664 |
| | CSF3 – CXCL1 | 0.5972 | CSF3 – LIF | 0.5245 |
| | IL-10 – CXCL10 | 0.5446 | IL-9 – IL-17 | 0.4792 |
| | IL-13 – CCL5 | 0.5487 | IL-10 – CCL5 | 0.5188 |
| | IL-12p40 – IL-2 * | 0.599 | IL-12p40 – CCL11 | 0.6154 |
| | IL-2 – IL-17 | 0.584 | IL-12p40 – IL-10 | 0.5386 |
| | IL-2 – IL-9 | 0.6145 | IL-12p40 – IL-1b | 0.6182 |
| | IL-3 – CCL5 | 0.5103 | IL-12p40 – IL-2 * | 0.6929 |
| | IL-9 – CCL11 | 0.6159 | IL-12p40 – VEGF | 0.5619 |
| | IL-9 – VEGF | 0.593 | IL-13 – IL-15 | 0.6092 |
| | LIF – IL-12p70 | 0.5975 | IL-15 – CXCL2 | 0.5573 |
| | LIF – IL-3 | 0.6275 | IL-2 – IL-1a | 0.6447 |
| | LIF – TNFa | 0.6473 | IL-4 – CCL11 | 0.6438 |
| | | | IL-4 – IL-13 | 0.6351 |
| | | | IL-4 – IL-17 | 0.6241 |
| Serum | CCL3 - IL-9 | 0.5579 | CXCL10 - CCL11 * | 0.5989 |
| | CSF3 - CCL2 | 0.5994 | IL-12p70 - IL-10 | 0.5341 |
| | CSF3 - LIF | 0.5211 | IL-12p70 - IL-1b | 0.5152 |
| | CXCL1 - IL-13 | 0.5262 | IL-12p70 - IL-4 | 0.5119 |
| | CXCL1 - IL-6 | 0.5773 | IL-12p70 - LIF | 0.5253 |
| | CXCL10 - CCL11 * | 0.5958 | IL-15 - CCL3 | 0.5476 |
| | CXCL10 - CCL2 | 0.7297 | IL-15 - IL-12p40 | 0.5958 |
| | CXCL10 - CCL3 | 0.6515 | IL-1b - TNFa | 0.5403 |
| | CXCL10 - CCL5 | 0.5392 | IL-7 - LIF | 0.5141 |
| | CXCL10 - CXCL9 | 0.6067 | | |

**Supplemental Table 1. Individual cytokine network links with mutual information (MI) scores in the hippocampus (clear) and serum (grey) of males and females under resting saline control conditions.** *Cytokine links conserved across sexes.

| Males | | | | Females | | | |
|---|---|---|---|---|---|---|---|
| **Cytokine Links** | **MI Score** | **Cytokine Links** | **MI Score** | **Cytokine Links** | **MI Score** | **Cytokine Links** | **MI Score** |
| CCL11 - CCL5 | 0.4303 | IL15 - CCL2 | 0.4602 | CCL4 - CCL5 | 0.3837 | IL-1b – IL-3 | 0.4363 |
| CCL11 – IL-12p40 | 0.4452 | IL1b - IL13 | 0.3137 | CCL5 - VEGF | 0.3914 | IL-2 - CCL11 | 0.4208 |
| CCL11 – IL-15 * | 0.4398 | IL1b - IL1a | 0.3824 | CSF2 - CSF3 | 0.3109 | IL-3 - LIF | 0.4359 |
| CCL11 – IL-17 | 0.4589 | IL2 - CXCL1 | 0.3826 | CSF3 - CXCL1 | 0.3779 | IL-4 - CXCL1 | 0.3438 |
| CCL11 – IL-1a | 0.5272 | IL3 - IL9 | 0.3953 | CXCL1 - CCL4 | 0.4339 | IL-4 - CXCL9 | 0.2138 |
| CCL4 - VEGF | 0.4094 | IL3 - VEGF | 0.3926 | CXCL10 - CCL2 | 0.4227 | IL-4 - CXCL9 | 0.3840 |
| CSF2 - CXCL1 | 0.3754 | IL6 - LIF | 0.3977 | CXCL2 - CCL2 | 0.4231 | IL-5 – IL-7 | 0.4425 |
| CXCL10 - CCL3 | 0.4679 | IL9 - CCL2 | 0.3923 | CXCL9 - IL3 | 0.2345 | IL-6 - CSF3 | 0.4518 |
| CXCL10 - CSF1 | 0.4477 | INFg - IL12p70 | 0.3644 | IL10 - CCL3 | 0.4407 | IL-7 - CCL11 | 0.4473 |
| CXCL10 - CSF3 | 0.4595 | INFg - IL5 | 0.3177 | IL10 - IL13 | 0.2844 | IL-9 - CXCL10 | 0.3766 |
| CXCL10 - CXCL9 | 0.4441 | TNFa - CXCL1 | 0.3771 | IL10 - IL1a | 0.4509 | IL-9 – IL-17 | 0.3638 |
| CXCL10 – IL-2 | 0.4418 | TNFa – IL-13 * | 0.3657 | IL12p40 - CSF1 | 0.4410 | IL-9 – IL-7 | 0.3586 |
| CXCL10 – IL-4 | 0.4814 | | | IL12p40 - LIF | 0.4425 | IL-9 - LIF | 0.4495 |
| CXCL10 – IL-6 | 0.4499 | | | IL12p40 - VEGF | 0.3535 | INFg - CCL3 | 0.2790 |
| IL-10 - CCL2 | 0.4776 | | | IL13 - VEGF | 0.4421 | INFg - CSF2 | 0.4393 |
| IL-10 - CSF2 | 0.3984 | | | IL15 - CCL11 * | 0.3768 | INFg - CXCL10 | 0.3737 |
| IL-10 - CXCL2 | 0.4455 | | | IL15 - CCL5 | 0.3305 | LIF - CCL2 | 0.3872 |
| IL-10 – IL-7 | 0.5019 | | | IL15 - CSF2 | 0.3725 | TNFa - CCL3 | 0.2878 |
| IL-10 - INFg | 0.3916 | | | IL15 - CXCL9 | 0.2384 | TNFa - CCL4 | 0.4142 |
| IL-10 - LIF | 0.3917 | | | IL1a - IL12p70 | 0.4377 | TNFa – IL-13 * | 0.4066 |
| IL-13 - CCL4 | 0.3111 | | | IL1b - CSF1 | 0.4326 | TNFa – IL-2 | 0.3658 |

**Supplemental Table 2. Individual cytokine network links with mutual information (MI) scores in the hippocampus of males and females 2 hours after LPS injection.** No networks were detected in the serum at 2 hours. *Cytokine links conserved across sexes.

| | Males | | | | Females | | | |
|---|---|---|---|---|---|---|---|---|
| | Cytokine Links | MI Score | Cytokine Links | MI Score | Cytokine Links | MI Score | Cytokine Links | MI Score |
| Hippocampus | CCL4 – IL-15 | 0.4986 | IL-1a – IL-12p40 | 0.4549 | CCL11 – VEGF* | 0.3063 | IL-1a - CCL3 | 0.4428 |
| | CCL4 – IL-7 | 0.4566 | IL-1b – IL-10 | 0.4106 | CCL2 - CCL5* | 0.4334 | IL-1b - CCL5 | 0.4220 |
| | CCL4 - INFg | 0.3808 | IL-2 – IL-1a* | 0.4533 | CCL4 - VEGF | 0.3527 | IL-2 – IL-1a* | 0.4603 |
| | CCL5 - CCL2* | 0.2787 | IL-3 - CCL11 | 0.3859 | CCL5 - CCL11 | 0.3264 | IL-2 – IL-7 | 0.4303 |
| | CCL5 - CSF1 | 0.3236 | IL-3 – LIF | 0.4672 | CSF2 - CCL4 | 0.4103 | IL-4 – IL-3 | 0.5229 |
| | CSF1 - CSF2 | 0.4090 | IL-4 - CSF2 | 0.4312 | CSF2 – IL-17 | 0.4341 | IL-4 – IL-5 | 0.3981 |
| | CSF3 - CXCL1* | 0.4007 | IL-4 – IL-12p70 | 0.4656 | CSF3 - CXCL1* | 0.5109 | IL-5 – IL-12p70 | 0.3026 |
| | CSF3 – IL-12p40 | 0.3548 | IL-4 – LIF | 0.4818 | CSF3 – IL-17 | 0.4349 | IL-6 – IL-12p70 | 0.4180 |
| | CXCL1 - CXCL2 | 0.3971 | IL-6 - CCL5 | 0.2354 | CXCL10 - VEGF | 0.2287 | IL-6 – LIF | 0.4882 |
| | CXCL1 – IL-5 | 0.3182 | IL-6 – IL-13 | 0.2946 | CXCL2 - CCL2 | 0.4056 | IL-7 – IL-12p40 | 0.4308 |
| | CXCL10 - CSF1 | 0.4112 | IL-6 – IL-7 | 0.2668 | CXCL2 – IL-1a | 0.4399 | IL-9 - CSF1 | 0.5231 |
| | CXCL10 - CXCL2 | 0.4525 | IL-7 - CCL3 | 0.2554 | CXCL9 - CXCL10 | 0.2559 | IL-9 - CXCL9 | 0.3899 |
| | CXCL9 - CSF3 | 0.4485 | IL-9 - CCL2 | 0.3302 | CXCL9 – IL-17 | 0.3928 | IL-9 - INFg | 0.4378 |
| | CXCL9 - CXCL2 | 0.4322 | LIF - CCL2 | 0.2996 | IL-10 – IL-7 | 0.4359 | IL-9 - LIF | 0.5180 |
| | IL-10 – IL-2 | 0.3729 | TNFa – IL-12p70 | 0.4303 | IL-12p40 - CCL4 | 0.4048 | IL-9 - TNFa | 0.5334 |
| | IL-10 – IL-9 | 0.4245 | TNFa – IL-15 | 0.4803 | IL-13 - CXCL9 | 0.4261 | IL-9 -IL-1b | 0.4675 |
| | IL-12p40 - CCL3 | 0.4153 | VEGF - CCL11* | 0.4357 | IL-13 – IL-2 | 0.4448 | INFg – IL-3 | 0.3526 |
| | IL-13 - CCL3 | 0.4060 | VEGF - INFg | 0.4413 | IL-15 - CSF1 | 0.4402 | INFg - VEGF | 0.3678 |
| | IL-17 - LIF | 0.4750 | VEGF – IL-1b | 0.4468 | IL-15 – IL-12p40 | 0.4359 | | |
| Serum | CCL2 - IL-12p40 | 0.3377 | IL-10 - CCL2 | 0.3835 | CCL2 - IL-13 | 0.4744 | CXCL2 - IL-13 | 0.5427 |
| | CCL2 - IL-13 | 0.2877 | IL-12p40 - IL-12p70 | 0.3885 | CCL2 - VEGF | 0.3964 | CXCL2 - LIF | 0.5408 |
| | CCL3 - CCL5 | 0.4119 | IL-15 - IL-17 | 0.4521 | CCL3 - CXCL2 | 0.4992 | IL-10 - CXCL9 | 0.2795 |
| | CCL4 - CCL11 | 0.4186 | IL-3 - IL-12p70 | 0.4425 | CCL3 - IL-7 | 0.2752 | IL-10 - IL-12p40 | 0.3389 |
| | CCL4 - CXCL1 | 0.4074 | IL-3 - IL-15 | 0.4116 | CCL4 - CCL5 | 0.3046 | IL-13 - CCL11 | 0.4764 |
| | CCL4 - VEGF | 0.4955 | IL-3 - IL-1b | 0.4487 | CCL4 - IL-1a | 0.3472 | IL-13 - CXCL1 | 0.4735 |
| | CCL5 - VEGF | 0.4342 | IL-3 - INFg | 0.4071 | CCL4 - IL-2 | 0.4732 | IL-1a - IL-7 | 0.2398 |
| | CSF1 - IL-17 | 0.4759 | IL-4 - IL-17 | 0.4412 | CCL4 - IL-4 | 0.4343 | IL-1b - LIF | 0.4683 |
| | CSF1 - IL-9 | 0.4420 | IL-5 - CSF3 | 0.3158 | CSF1 - IL-2 | 0.3659 | IL-2 - IL-3 | 0.4538 |
| | CSF1 - LIF | 0.5059 | IL-5 - CXCL1 | 0.3420 | CSF1 - IL-7 | 0.3224 | IL-3 - IL-12p70 | 0.4574 |
| | CSF2 - IL-4 | 0.4491 | IL-5 - IL-10 | 0.3533 | CSF2 - IL-9 | 0.3912 | IL-4 - IL-12p70 | 0.4458 |
| | CSF3 - INFg * | 0.2495 | IL-5 - IL-6 | 0.2953 | CSF3 - IL-15 | 0.4252 | IL-5 - LIF | 0.4509 |
| | CXCL10 - CXCL2 | 0.4369 | IL-6 - CCL11 | 0.2423 | CSF3 - IL-17 | 0.4767 | IL-6 - CXCL10 | 0.3561 |
| | CXCL2 - IL-2 | 0.4365 | IL-6 - IL-12p70 | 0.2742 | CSF3 - IL-3 | 0.4635 | IL-6 - IL-12p40 | 0.3505 |
| | CXCL9 - CSF1 | 0.5321 | IL-7 - LIF | 0.4586 | CSF3 - IL-9 | 0.4121 | IL-9 - IL-12p40 | 0.3804 |
| | CXCL9 - CXCL2 | 0.4600 | TNFa - CCL3 | 0.4256 | CSF3 - INFg * | 0.4823 | LIF - CXCL9 | 0.4579 |
| | CXCL9 - IL-13 | 0.4915 | TNFa - IL-1a | 0.3047 | CSF3 - VEGF | 0.4693 | TNFa - CXCL2 | 0.4799 |
| | CXCL9 - VEGF | 0.5342 | TNFa - IL-9 | 0.4252 | CXCL10 - IL-5 | 0.4457 | | |

**Supplemental Table 3. Individual cytokine network links with mutual information (MI) scores in the hippocampus (white) and serum (grey) of males and females 6 hours after LPS injection.** *Cytokine links conserved across sexes.

| | Males | | | | Females | | | |
|---|---|---|---|---|---|---|---|---|
| | Cytokine Links | MI Score | Cytokine Links | MI Score | Cytokine Links | MI Score | Cytokine Links | MI Score |
| Hippocampus | CCL2 - CCL5 | 0.4433 | IL-2 - CCL3 | 0.3975 | CCL3 - CCL5 | 0.2941 | IL-3 – IL-17 | 0.4645 |
| | CSF1 - VEGF | 0.4168 | IL-2 – IL-17 | 0.3747 | CSF1 - CXCL1 | 0.3285 | IL-3 – IL-7 * | 0.4441 |
| | CSF2 - CCL4 * | 0.3958 | IL-2 – IL-1a | 0.3657 | CSF2 - CCL2 | 0.4165 | IL-3 - INFg | 0.4660 |
| | CSF2 - CSF3 | 0.4744 | IL-3 - CSF1 | 0.4441 | CSF2 - CCL4 * | 0.4194 | IL-3 - VEGF | 0.4852 |
| | CSF3 - CCL3 | 0.3885 | IL-3 – IL-15 * | 0.4306 | CSF2 – LIF | 0.4272 | IL-5 – LIF | 0.5201 |
| | CSF3 - CXCL1 | 0.4941 | IL-3 – IL-7 * | 0.3744 | CSF3 - CCL11 | 0.2079 | IL-6 - CCL2 | 0.3841 |
| | CSF3 – IL-5 | 0.4281 | IL-3 – LIF | 0.3123 | CSF3 - CCL2 | 0.3993 | IL-6 - CSF1 | 0.3300 |
| | CXCL10 – IL-12p70 | 0.3820 | IL-4 - CCL11 | 0.3403 | CXCL1 - CCL11 | 0.3800 | IL-6 – IL-7 * | 0.4298 |
| | CXCL2 - CCL2 | 0.4665 | IL-4 - CSF1 | 0.4515 | CXCL1 - CCL5 | 0.3551 | IL-7 - CCL3 | 0.4327 |
| | CXCL2 – IL-5 | 0.4350 | IL-6 - CCL4 | 0.3930 | CXCL2 - LIF | 0.5065 | IL-9 – IL-12p40 * | 0.4362 |
| | CXCL9 - CXCL1* | 0.4818 | IL-6 – IL-7 * | 0.3611 | CXCL9 - CXCL1* | 0.3595 | TNFa - CXCL2 | 0.3750 |
| | CXCL9 – IL-12p70 | 0.4078 | IL-9 - CCL11 | 0.4772 | CXCL9 - CXCL10 | 0.4004 | TNFa – IL-12p70 | 0.4424 |
| | IL-10 - CSF1 | 0.3444 | IL-9 – IL-12p40 * | 0.5041 | IL-1a – IL-12p40 | 0.4199 | | |
| | IL-10 - CXCL2 | 0.4022 | INFg - CCL3 | 0.2976 | IL-1b – IL-12p40 | 0.4200 | | |
| | IL-10 - IL17 | 0.3879 | INFg - CXCL10 | 0.2269 | IL-1b – IL-2 | 0.4181 | | |
| | IL-12p70 - LIF | 0.3280 | INFg - VEGF | 0.3258 | IL-2 – IL-7 | 0.4290 | | |
| | IL-13 - CCL2 | 0.3973 | LIF - CCL2 | 0.3440 | IL-3 – IL-10 | 0.4696 | | |
| | IL-13 - CXCL10 | 0.3058 | TNFa - CCL4 | 0.4100 | IL-3 – IL-12p40 | 0.4445 | | |
| | IL-13 – IL-9 | 0.4473 | TNFa - CXCL1 | 0.3971 | IL-3 – IL-12p70 | 0.4408 | | |
| | IL-1a – IL-9 | 0.4502 | TNFa – IL-3 | 0.4674 | IL-3 – IL-13 | 0.4479 | | |
| | IL-1b – IL-4 | 0.4450 | | | IL-3 – IL-15 * | 0.4455 | | |
| Serum | CCL2 - CXCL2 | 0.4001 | IL-1b - IL-4 | 0.4287 | CCL11 - CXCL2 | 0.3860 | IL-2 - IL-9 * | 0.4596 |
| | CCL4 - VEGF | 0.3671 | IL-1b - TNFa | 0.3597 | CCL11 - IL-1a | 0.3739 | IL-4 - IL-12p40 | 0.4417 |
| | CCL5 - CSF3 | 0.3507 | IL-5 - CCL11 * | 0.2958 | CCL11 - IL-5 * | 0.4090 | IL-4 - IL-9 | 0.3903 |
| | CCL5 - CXCL10 | 0.3757 | IL-5 - LIF * | 0.4304 | CCL4 - CCL5 | 0.3786 | IL-4 - VEGF | 0.4522 |
| | CCL5 - CXCL2 | 0.4669 | IL-6 - CCL2 | 0.4158 | CCL5 - IL-1a | 0.3802 | IL-5 - LIF * | 0.4314 |
| | CSF1 - IL-4 | 0.3845 | IL-6 - IL-10 | 0.4142 | CSF1 - CSF3 | 0.4585 | IL-6 - CSF1 | 0.4612 |
| | CSF1 - IL-7 | 0.2672 | IL-7 - LIF | 0.3103 | CSF1 - VEGF | 0.4644 | IL-7 - CXCL9 | 0.5414 |
| | CXCL10 - IL-12p40 | 0.3635 | IL-9 - CCL3 | 0.4946 | CSF2 - IL-3 | 0.4074 | IL-7 - IL-13 | 0.5257 |
| | CXCL2 - IL-5 | 0.3225 | IL-9 - CCL4 | 0.4900 | IL-10 - CCL2 | 0.4773 | IL-7 - IL-15 * | 0.5164 |
| | CXCL9 - CCL2 | 0.3909 | IL-9 - CXCL1 | 0.4924 | IL-10 - CCL3 | 0.4663 | IL-7 - IL-2 | 0.5010 |
| | CXCL9 - CXCL10 | 0.3427 | IL-9 - IL-10 | 0.4538 | IL-10 - CCL4 | 0.3862 | IL-7 - IL-3 | 0.5248 |
| | CXCL9 - IL-1a | 0.3138 | IL-9 - IL-12p70 | 0.4757 | IL-10 - CXCL1 | 0.3706 | IL-7 - INFg | 0.5342 |
| | CXCL9 - TNFa | 0.3961 | IL-9 - IL-13 | 0.4953 | IL-12p40 - IL-12p70 | 0.4112 | TNFa - IL-12p70 | 0.4251 |
| | IL-15 - CSF2 | 0.4174 | IL-9 - IL-15 | 0.4255 | IL-17 - CXCL9 | 0.4666 | TNFa - INFg | 0.4542 |
| | IL-15 - IL-12p40 | 0.2547 | IL-9 - IL-17 | 0.4909 | IL-17 - LIF | 0.4286 | IL-2 - IL-9 * | 0.4596 |
| | IL-15 - IL-7 * | 0.3516 | IL-9 - IL-2 * | 0.4958 | IL-1b - CCL2 | 0.4667 | IL-4 - IL-12p40 | 0.4417 |
| | IL-1a - VEGF | 0.4396 | IL-9 - IL-3 | 0.4919 | IL-1b - CXCL10 | 0.4501 | IL-4 - IL-9 | 0.3903 |
| | IL-1b - CSF3 | 0.4280 | IL-9 - INFg | 0.4921 | IL-1b - IL-3 | 0.4672 | IL-4 - VEGF | 0.4522 |

**Supplemental Table 4. Individual cytokine network links with mutual information (MI) scores in the hippocampus (white) and serum (grey) of males and females 24 hours after LPS injection.**
*Cytokine links conserved across sexes.

| | Males | | | | Females | | | |
|---|---|---|---|---|---|---|---|---|
| | **Cytokine Links** | **MI Score** | **Cytokine Links** | **MI Score** | **Cytokine Links** | **MI Score** | **Cytokine Links** | **MI Score** |
| Hippocampus | CSF1 - CCL2 | 0.4540 | IL-6 – IL-15 | 0.4587 | CSF1 – IL-12p40 | 0.4945 | IL-2 - CCL3 | 0.4422 |
| | CSF1 – IL-9 | 0.4862 | IL-7 - CCL2 | 0.5057 | CSF1 – IL-7 | 0.4747 | IL-3 - CCL3 | 0.5145 |
| | CSF2 – LIF | 0.3167 | IL-7 - CCL5 | 0.5172 | CSF3 – IL-3 | 0.4309 | IL-3 – IL-9 | 0.5312 |
| | CXCL1 – IL-9 | 0.4982 | IL-7 - CSF3 | 0.5219 | CXCL9 - CXCL1 | 0.4699 | IL-6 - CCL2 | 0.4165 |
| | CXCL10 - CSF2 | 0.3231 | IL-7 - CXCL9 | 0.6315 | CXCL9 – IL-3 | 0.5636 | IL-6 - CXLC2 | 0.4437 |
| | IL-10 - CXLC2 | 0.4845 | IL-7 – IL-10 | 0.5209 | IL-10 – IL-7 | 0.4850 | IL-6 – IL-3 | 0.4138 |
| | IL-10 – IL-12p70 | 0.4924 | IL-7 – IL-12p40 | 0.5920 | IL-13 - CXCL1 | 0.4027 | IL-6 - INFg | 0.4086 |
| | IL-12p40 - CCL3 | 0.5390 | IL-7 – IL-1a | 0.5468 | IL-13 – IL-7 * | 0.4981 | IL-6 - VEGF | 0.4071 |
| | IL-12p40 - VEGF | 0.5091 | IL-7 – IL-3 * | 0.5283 | IL-15 - CXLC10 | 0.4964 | INFg - CCL4 | 0.5135 |
| | IL-13 - CXCL10 | 0.4938 | IL-7 – IL-6 | 0.5191 | IL-15 – IL-3 | 0.5314 | INFg - CSF3 | 0.4466 |
| | IL-13 - VEGF | 0.4913 | IL-7 - INFg | 0.4333 | IL-1b - CCL11 | 0.5712 | | |
| | IL-1b – IL-12p40 * | 0.5187 | IL-7 – LIF | 0.5356 | IL-1b - CCL5 | 0.6003 | | |
| | IL-1b – IL-2 * | 0.4852 | IL-7 - TNFa | 0.4623 | IL-1b - CSF2 | 0.6118 | | |
| | IL-2 - CXCL1 | 0.5143 | IL-9 - CCL11 | 0.3962 | IL-1b - CXCL9 | 0.6911 | | |
| | IL-2 - CXCL10 | 0.4958 | IL-9 - CCL4 | 0.3855 | IL-1b – IL-12p40 * | 0.6138 | | |
| | IL-2 - VEGF | 0.5025 | IL-9 – IL-12p40 | 0.6190 | IL-1b – IL-12p70 | 0.5331 | | |
| | IL-3 – IL-5 | 0.4702 | IL-9 – IL-17 | 0.5395 | IL-1b – IL-1a | 0.6229 | | |
| | IL-4 - CCL11 | 0.3626 | TNFa - CXLC2 | 0.4654 | IL-1b – IL-2 * | 0.5198 | | |
| | IL-4 - CSF2 | 0.3108 | IL-7 - CCL2 | 0.5057 | IL-1b - LIF | 0.6399 | | |
| | IL-4 – IL-12p70 | 0.3636 | | | IL-1b - TNFa | 0.4897 | | |
| Serum | CCL2 - CSF2 | 0.4613 | IL-12p70 - CXCL1 | 0.3025 | IL-1b - CCL4 | 0.4644 | CSF1 - INFg | 0.7173 |
| | CCL2 - CXCL9 | 0.2634 | IL-15 - CCL3 | 0.4458 | IL-1b - CSF1 | 0.5190 | CSF1 - IL-1a | 0.6324 |
| | CCL3 - IL-13 | 0.4031 | IL-15 - IL-10 | 0.4089 | TNFa - IL-9 | 0.5037 | CSF1 - IL-2 | 0.6521 |
| | CCL3 - IL-5 | 0.4782 | IL-15 - IL-12p40 | 0.4411 | TNFa - VEGF | 0.4313 | CSF1 - IL-7 * | 0.7118 |
| | CCL4 - CCL11 | 0.4341 | IL-17 - CCL4 | 0.4325 | IL-6 - CCL2 | 0.5678 | CSF1 - IL-12p40 | 0.5639 |
| | CCL4 - VEGF * | 0.4154 | IL-17 - CCL5 | 0.3757 | IL-6 - CSF3 | 0.6032 | CSF1 - CXCL1 | 0.7890 |
| | CCL5 - INFg | 0.4124 | IL-17 - IL-4 | 0.4525 | IL-10 - CSF1 | 0.4770 | CSF2 - IL-12p40 | 0.4240 |
| | CSF1 - IL-7 * | 0.3891 | IL-17 - TNFa | 0.3837 | IL-10 - IL-12p70 | 0.4268 | CSF2 - CXCL9 | 0.4912 |
| | CSF1 - VEGF | 0.3346 | IL-1b - IL-13 | 0.3854 | IL-10 - CXCL9 | 0.3991 | CSF2 - VEGF | 0.3790 |
| | CSF2 - IL-3 | 0.3722 | IL-1b - IL-17 | 0.4661 | CCL3 - CSF1 | 0.5857 | CSF3 - CXCL1 | 0.6517 |
| | CSF3 - IL-3 | 0.3590 | IL-2 - IL-9 | 0.4017 | CCL4 - CSF3 | 0.4974 | IL-4 - IL-12p70 | 0.2935 |
| | CSF3 - IL-7 | 0.4101 | IL-3 - CXCL1 | 0.3288 | CCL4 – VEGF * | 0.4730 | IL-9 - IL-12p70 | 0.4687 |
| | CSF3 - LIF | 0.3561 | IL-4 - IL-1a | 0.4763 | CCL5 - CXCL2 | 0.5758 | IL-9 - IL-15 | 0.4602 |
| | CXCL10 - IL-4 | 0.4859 | IL-5 - IL-9 | 0.4667 | CCL5 - CXCL10 | 0.6628 | IL-9 - IL-17 | 0.4764 |
| | CXCL2 - IL-15 | 0.4452 | IL-6 - CXCL9 | 0.4015 | CCL5 - CCL11 | 0.5737 | IL-9 - CXCL1 | 0.7121 |
| | CXCL2 - IL-2 | 0.4238 | IL-6 - IL-12p70 | 0.3764 | CXCL2 - IL-9 | 0.6607 | IL-13 - IL-17 | 0.4572 |
| | CXCL2 - IL-5 | 0.4355 | IL-7 - IL-12p40 | 0.3689 | CXCL10 - IL-1a | 0.9085 | IL-13 - CXCL9 | 0.4517 |
| | CXCL2 - IL-6 | 0.4119 | INFg - CXCL1 | 0.3531 | CXCL10 - IL-5 | 0.6163 | | |
| | IL-10 - LIF | 0.4106 | TNFa - CCL2 | 0.3717 | CXCL10 - CXCL9 | 0.5611 | | |

**Supplemental Table 5. Individual cytokine network links with mutual information (MI) scores in the hippocampus (white) and serum (grey) of males and females 48 hours after LPS injection.** *Cytokine links conserved across sexes.

| | Males | | | | Females | | | |
|---|---|---|---|---|---|---|---|---|
| | **Cytokine Links** | **MI Score** | **Cytokine Links** | **MI Score** | **Cytokine Links** | **MI Score** | **Cytokine Links** | **MI Score** |
| Hippocampus | CCL3 - VEGF | 0.4473 | IL-1b – IL-12p40 | 0.4751 | CCL4 - CCL5 | 0.5446 | IL-15 – IL-5 | 0.4705 |
| | CSF1 - CXCL2 | 0.5464 | IL-1b – IL-3 | 0.6593 | CSF2 – IL-9 | 0.5411 | IL-1a - CCL4 | 0.5661 |
| | CSF2 - CCL5 | 0.4774 | IL-3 – IL-12p70 * | 0.5752 | CSF3 – IL-9 | 0.5323 | IL-1b - CCL11 | 0.7447 |
| | CSF3 - CCL5 | 0.6989 | IL-4 - CSF3 | 0.6074 | CXCL1 – IL-12p40 | 0.4474 | IL-1b - CCL5 | 0.7246 |
| | CSF3 – IL-3 | 0.7570 | IL-4 - CXCL2 | 0.4629 | CXCL2 - CCL4 | 0.4861 | IL-1b - CXCL9 | 0.7431 |
| | CSF3 - INFg | 0.5954 | IL-4 – IL-5 | 0.4826 | CXCL9 - CXCL1 * | 0.5111 | IL-1b - TNFa | 0.5711 |
| | CSF3 - LIF | 0.4091 | IL-5 - CCL2 | 0.4831 | CXCL9 - CXCL10 | 0.5054 | IL-2 – IL-1a | 0.4289 |
| | CXCL10 - VEGF | 0.4365 | IL-6 – IL-15 | 0.4782 | IL-10 - CXCL9 | 0.4867 | IL-5 - CCL5 | 0.4381 |
| | CXCL9 - CXCL1 * | 0.4827 | IL-7 - CCL11 | 0.5647 | IL-10 – IL-7 | 0.4715 | IL-6 – IL-4 | 0.4576 |
| | CXCL9 – IL-12p40 | 0.4366 | IL-7 - CCL5 | 0.6948 | IL-10 – VEGF * | 0.4604 | IL-9 - CCL5 | 0.5892 |
| | CXCL9 – IL-1a | 0.4463 | IL-7 - CXCL1 | 0.4832 | IL-12p70 - CCL3 | 0.5163 | IL-9 – IL-2 | 0.4557 |
| | IL-10 - CSF2 | 0.5182 | IL-7 - CXCL10 | 0.5720 | IL-12p70 - CCL4 | 0.6280 | IL-9 - LIF | 0.5849 |
| | IL-10 – IL-13 | 0.5410 | IL-7 – IL-3 | 0.6913 | IL-12p70 - CSF1 | 0.4946 | LIF - CCL2 | 0.4205 |
| | IL-10 – IL-17 | 0.5145 | IL-7 – IL-6 | 0.4918 | IL-12p70 – IL-1b | 0.7811 | | |
| | IL-10 – IL-1a | 0.4693 | IL-9 – IL-17 | 0.4689 | IL-12p70 – IL-3 * | 0.6324 | | |
| | IL-10 – IL-2 | 0.5207 | INFg - CCL4 | 0.5621 | IL-12p70 – IL-4 | 0.5937 | | |
| | IL-10 – VEGF * | 0.5569 | TNFa - CCL5 | 0.6106 | IL-12p70 - INFg | 0.5094 | | |
| | IL-15 - CXCL2 | 0.4812 | TNFa – IL-5 | 0.4973 | IL-12p70 - LIF | 0.5224 | | |
| | IL-15 – IL-1a | 0.4712 | | | IL-13 - CCL5 | 0.5121 | | |
| Serum | CCL11 - CCL2 | 0.5381 | IL-13 - IL-7 | 0.4409 | CCL2 - CSF1 | 0.6669 | CXCL1 - CCL11 * | 0.6044 |
| | CCL11 - CCL4 | 0.5143 | IL-17 - CCL5 | 0.4731 | CCL2 - CSF3 | 0.4415 | CXCL1 - CCL5 | 0.5824 |
| | CCL11 - CXCL1 * | 0.5548 | IL-17 - IL-12p70 | 0.4758 | CCL2 - CXCL9 | 0.5891 | CXCL1 - CSF2 | 0.7013 |
| | CCL11 - CXCL2 | 0.4943 | IL-17 - IL-15 | 0.4788 | CCL2 - IL-15 | 0.6751 | CXCL1 - IL-12p40 | 0.5956 |
| | CCL11 - IL-10 | 0.5224 | IL-17 - VEGF | 0.4868 | CCL2 - IL-6 | 0.4967 | CXCL1 - IL-17 | 0.5142 |
| | CCL11 - IL-17 | 0.5372 | IL-1a - IL-9 | 0.4478 | CCL2 - TNFa | 0.6032 | CXCL2 - IL-3 | 0.3845 |
| | CCL11 - IL-5 | 0.5533 | IL-1b - IL-2 | 0.4510 | CCL3 - IL-17 | 0.4257 | IL-10 - CSF2 | 0.5295 |
| | CCL3 - IL-9 | 0.3151 | IL-1b - IL-3 | 0.4506 | CCL4 - CCL2 | 0.4958 | IL-10 - VEGF | 0.4383 |
| | CCL3 - LIF | 0.1356 | IL-3 - IL-12p70 | 0.4620 | CCL4 - CSF2 | 0.7944 | IL-12p70 - VEGF | 0.4326 |
| | CSF1 - IL-5 | 0.2650 | IL-4 - INFg * | 0.4543 | CCL4 - CXCL10 | 0.5520 | IL-1a - CXCL9 | 0.5488 |
| | CSF1 - LIF | 0.2546 | IL-6 - IL-1a | 0.4589 | CCL4 - IL-1b | 0.6146 | IL-3 - VEGF | 0.4164 |
| | CSF2 - CXCL9 | 0.3478 | IL-6 - INFg | 0.4522 | CCL4 - IL-2 | 0.3933 | IL-9 - CXCL9 | 0.4589 |
| | CXCL10 - CSF3 | 0.4417 | TNFa - IL-10 | 0.4662 | CCL4 - INFg | 0.5125 | INFg - IL-12p40 | 0.5112 |
| | CXCL10 - IL-12p40 | 0.4602 | | | CCL4 - VEGF | 0.4546 | INFg - IL-1a | 0.5619 |
| | CXCL2 - CSF3 | 0.4134 | | | CCL5 - CXCL2 | 0.5174 | INFg - IL-4 * | 0.5316 |
| | IL-12p40 - CXCL9 | 0.4478 | | | CSF2 - IL-13 | 0.5802 | INFg - LIF | 0.4300 |
| | IL-13 - IL-2 | 0.4435 | | | CSF2 - IL-5 | 0.5083 | TNFa - CXCL2 | 0.5321 |
| | IL-13 - IL-4 | 0.4535 | | | CSF3 - IL-5 | 0.4570 | TNFa - IL-7 | 0.5748 |

**Supplemental Table 6. Individual cytokine network links with mutual information (MI) scores in the hippocampus (white) and serum (grey) of males and females 168 hours after LPS injection.** *Cytokine links conserved across sexes.
